# Leveraging structurally-informed bioorthogonal conjugation for alpha synuclein fibril surveillance

**DOI:** 10.1101/2025.09.12.675751

**Authors:** Rebecca A. Jenkins, Samantha Wu, Cameron W. Flowers, Lily J. Taylor, Gretchen Fujimura, Andrés Heredia, Chuanqi Sun, Michael R. Sawaya, Joseph A. Loo, Jose A. Rodriguez

**Affiliations:** Department of Chemistry and Biochemistry; UCLA-DOE Institute for Genomics and Proteomics; STROBE, NSF Science and Technology Center; University of California, Los Angeles (UCLA); Los Angeles, CA 90095, USA; Department of Chemistry and Biochemistry; UCLA-DOE Institute for Genomics and Proteomics; University of California, Los Angeles (UCLA); Los Angeles, CA 90095, USA

**Keywords:** Synuclein, amyloid, bioorthogonal, fluorescence, fibril

## Abstract

Alpha synuclein (α-syn) amyloid fibrils are etiological agents of neurodegenerative disease. To better understand the molecular and cellular basis for their persistence and spread, we implemented a fluorophore labeling strategy to track pre-formed recombinant α-syn fibrils in solution and in cells. We leveraged amber codon mediated incorporation of a tetrazine-based artificial amino acid (TetV2.0) to install a strained trans-cyclooctene (sTCO)-conjugated Janeliafluor (JF549), at four structurally-informed sites on human α-syn: residues 4, 60, 96 and 136. A structure of fibrils labeled at residue 60 showed density consistent with TetV2.0 incorporation and a core structure similar to recombinant type 1M α-syn fibrils. Coupling of TetV2-containing fibrils to sTCO-JF549 proceeded under mild buffer conditions and in the presence of the disease-associated cofactor and cytotoxic lipid, psychosine. Labeled psychosine-induced fibrils retained their polymorph-specific features during labeling; their structure resembled that of type 1A recombinant α-syn fibrils and showed a solvent exposed site at residue 60, amenable to bioorthogonal conjugation. Labeled fibrils also seeded new fibril growth *in vitro*, and induced the formation of puncta in α-syn FRET biosensor HEK293T cells, facilitating the simultaneous tracking of exogenous and endogenous α-syn aggregates and their localization within the cells. This approach opens a correlative bridge between fibril structure and function in living systems.

## Introduction

The presence of α-synuclein (α-syn) fibrils is correlated with pathology in various neurodegenerative diseases including Parkinson’s, dementia with Lewy bodies (DLB), Parkinson’s disease dementia (PDD) and multiple system atrophy (MSA)^1,2^. The structures of recombinant and tissue-derived α-syn fibrils have been determined by cryoEM and exhibit distinct folds^3,4^. Many of these fibrils share structural motifs and can be organized into polymorph classes that appear characteristic of a disease type or recombinant growth condition^5^. Notably, *in vitro* grown species appear distinct from disease-associated, tissue-derived polymorphs, which have been a challenge to recombinantly replicate^6^.

The spread of α-syn fibrils between cells and tissues has been attributed to prion-like behavior^7^. The tracking of endogenous or exogenous aggregates promises to help our understanding of α-syn fibril persistence, templating and spread^8^. The templating or seeding of recombinant, monomeric α-syn by pre-formed fibrils has been demonstrated under various conditions *in vitro*, tracked via thioflavin fluorescence^9^, and in cellular biosensor models via CFP and YFP-tagged α-syn that yield a fluorescence resonance energy transfer (FRET) signal upon aggregation^10^. These biosensor cell lines are increasingly used as tools to study seeding activity from various oligomeric and fibril species and show potential in the classification of distinct α-syn fibril strains or polymorphs^11^. However, the mechanism by which these fibrils propagate or seed within cells and the parameters that influence specific fibril polymorph structure persistence and faithful templating remain unclear.

Fibrils of α-syn have also been associated with the lysosomal storage disorders, including Krabbe disease (KD), where pathology is linked to the toxic buildup of the glycolipid psychosine^12^. In KD, the neuronal accumulation of psychosine also results in axon demyelination and neurodegeneration^13^. In addition, potentially seed competent α-syn inclusions have been identified in brains of KD mouse models and human patients^14,15^. However, a structure of α-syn fibrils from KD tissues is lacking and, while α-syn fibrils have been grown *in vitro* in the presence of psychosine, their structure, their capacity for templated spread, and their toxicity to cell models remain unknown.

The covalent chemical conjugation of fluorophores to amyloid fibrils can suffer from limited specificity, as occurs with NHS ester-based conjugation, require pH alterations^16^, or the incorporation of free, solvent accessible thiols into the primary sequence ^17^. Despite its widespread use, maleimide-thiol chemistry can result in undesired products^18^ and disruption of native disulfides^19^ for general protein labeling considerations. For α-syn fibrils specifically, maleimide-thiol chemistry has been used extensively to create site-specific labeled fibrils with maleimide containing fluorophores such as BODIPY^20^, Hilyte Fluor 488^21^ and Alexa Fluor 488^22^, where labeled monomer is fibrillized and labeled fibrils tracked in cells to understand the molecular basis for templated seeding and cellular uptake. Here, we implement a different type of site-specific labeling by leveraging chemical biology approaches to effectively place fluorescent labels on pre-formed amyloid fibrils. We chose to label pre-formed fibrils over monomers to minimize potential influence of fluorophores on fibril morphology^23^. This approach is attractive given its compatibility with diverse artificial amino acids, its site-selectivity without the need to disrupt native cysteines, and its bioorthogonality^24^. We apply our methodology to α-syn fibrils, including those formed in the presence of psychosine, as a model for potential fibril folds found in KD. We use strained cyclooctene-mediated bioorthogonal chemistry to install established fluorophores onto tetrazine-containing artificial amino acids at select sites on the α-syn sequence by genetic code expansion. Doing this on pre-formed fibrils offers a facile way of generating and tracking them, as they retain their initial structure and can be used in amyloid aggregation cell models such as biosensors. Conjugation of pre-formed fibrils permits their structural analysis. Installment of TetV2.0 at α-syn residue 60 results in a core fibril structure that, as revealed by cryoEM, is similar to that of the type 1M α-syn fibril strain. The mild bioorthogonal conjugation conditions are compatible with psychosine-induced α-syn fibrils, whose cryoEM structure reveals a similarity to the type 1A recombinant α-syn fibril polymorph and the ‘strain B’ α-syn fibril structure^25^. These results collectively demonstrate the utility of fluorophore bioconjugation strategies for tracking recombinant α-syn fibril polymorphs in disease models, including in biosensor cells.

## Results

### Characterization of fluorophore conjugated, pre-formed a-syn fibrils

We first evaluated the impact of amine-specific covalent labeling of pre-formed α-syn fibrils. ATTO-550 dye equipped for NHS-ester coupling was selected for this purpose, given its successful and robust use for visualizing α-syn fibrils in cells^26^. For this purpose, wild-type α-syn monomers were recombinantly expressed and purified from *E.coli* for aggregation studies (Figure S1). According to cryoEM structures of recombinant α-syn fibril cores, ATTO-550 could label multiple solvent exposed lysine residues (Figure S2A-C), assuming pre-formed fibrils were structurally unaffected by the conjugation procedure used to covalently link ATTO-550 to solvent-exposed lysines and or N-termini of fibril components (Figure S2B). Fibrils were labeled in an elevated pH solution at sub-stoichiometric ratios of dye to protein ranging from 0.1 to 0.5, to limit the potential steric interference between adjacent labeled sites along the fibril (Figure 2D-G).

SDS-PAGE of urea-denatured fibrils showed a fluorescent band at the expected molecular weight of monomeric α-syn, 14 kilodaltons (Figure S2C,F), and absorbance and emission spectra of monomer and fibril conjugates showed peaks similar to free dye (Figure S3). We characterized the morphology of wild-type α-syn fibrils self-assembled in solutions containing ionic additive and/or tris buffered saline (Figure S2D). Negative-stain electron microscopy (nsEM) suggested that overall fibril morphologies were minimally perturbed post labeling (Figure S2D), but that the high pH environment used for covalent attachment had the capacity to unravel fibrils over time (Figure S2E). This indicated that careful monitoring of fibril preparations would be required before and after NHS-ester-based dye conjugation. We also prepared ATTO-550 conjugated α-syn monomers under neutral and basic pH conditions at a sub-stoichiometric ratio of dye to protein (0.1-fold) and equal molar amounts of dye to protein. Under the same reaction conditions as ATTO-550 fibril conjugation, SDS-PAGE revealed a fluorescent band at the expected molecular weight of monomeric α-syn, 14 kilodaltons, for all conditions. The 1:1 dye to protein lanes had a wider band compared to the sub-stoichiometric conditions, but labeling at pH 7 versus pH 9.6 seemed to have no significant difference in band width (Figure S2F). Liquid chromatography mass spectrometry (LC-MS) of ATTO-550 conjugated α-syn monomers at excess dye to protein (10-fold) ratios revealed a heterogeneous pattern of conjugation that included multiple dyes conjugated to a single polypeptide, with populations of α-syn monomer coupled to 1-2 ATTO-550 molecules compared to its unlabeled counterpart (Figure S2G).

Clusters of ATTO-550-α-syn fibril seeds, evident by nsEM (Figure S4A), were readily observed by fluorescence within CellTrace™ Violet-labeled HEK293T cells after lipid-mediated transduction (Figure S4B). ATTO-550-α-syn fibril signal showed distinct puncta that colocalized with cell violet signal used as a stable fluorescent marker for intracellular proteins (Figure S4B). Similar ATTO-550-α-syn fibril puncta were observed in A53T α-syn biosensor cells^10^ (Figure S4C) and could be simultaneously imaged alongside FRET puncta representing endogenous, nascent α-syn aggregates. FRET puncta typically appeared ∼24h after α-syn fibril transduction, and upon visual inspection exhibited limited colocalization with ATTO-550-α-syn fibrils (Figure S4C), although that distribution may have been affected by the ATTO-550 labeling. It is possible then a population of unlabeled fibrils was able to seed, or that labeled fibrils were cleaved within the cell to yield a fibril fragment that seeded but did not carry the label.

### Selective, bioorthogonal conjugation of α-syn fibrils using noncanonical amino acids

To selectively and bioorthogonally label α-syn fibrils, we implemented an amber codon mediated site-specific incorporation of the noncanonical amino acid (nCAA), 1,2,4,5-tetrazine-ethyl (Tet2-Et), in *E.coli* using a non-native aminoacyl tRNA synthetase (aaRS)^27^. This resulted in α-syn monomers bearing a single Tet2-Et residue (Figure S5). We leveraged the reaction of Tet2-Et residues with strained trans cyclo-octene (sTCO)-containing compounds through an inverse-electron-demand Diels-Alder reaction, which occurs efficiently under mild buffer conditions^28^. Informed by the known structures of recombinant α-syn fibril polymorphs, we implemented the incorporation at four amino acid sites on α-syn (F4, K60, K96 and Y136) that might be suitable for Tet2-Et incorporation and dye coupling based on potential solvent accessibility and steric constraints (Figure S6A). K60 and K96 reside in the fibril core but are not buried within a protofilament or at the interface of the two protofilaments. In contrast, F4 and Y136 were chosen because they do not reside in the ordered fibril core based on cryoEM densities for α-syn fibrils, and are expected to be solvent exposed and amendable for Tet2-ethyl substitution and conjugation. Accordingly, a solid-state NMR structure of full-length α-syn fibrils shows these amino acids are located in disordered regions that flank an ordered core^29^. Tet2-Et was effectively incorporated into α-syn, allowing its expression and purification from *E.coli* with similar yield and purity to wild type (Figure S1, Figure S7). We used LC-MS to quantify the efficiency of Tet2-Et amino acid incorporation and did not detect any truncation products that would arise from premature termination of translation for F4, K60, K96 and Y136 α-syn monomers (Figure S8).

We investigated the impact of Tet2-Et incorporation on the kinetics of α-syn monomer aggregation, as monitored by Thioflavin T (ThT) incorporation (Figure 1A), and assessed the morphology of the assembled fibrils by nsEM (Figure 1B). We also evaluated the impact of sTCO-JF549 labeling on fibrillization and noted that K60 and Y136 monomers conjugated to JF549 retained the capacity to fibrillize. However, labeled K60-α-syn (K60^FL^-α-syn) monomers had a longer lag phase in their ThT curves compared to their unlabeled counterparts (Figure S9A), and nsEM analysis showed they tended to form amorphous aggregates rather than mature fibrils (Figure S9B). We also evaluated fibrillization by ATTO-550-labeled wild-type α-syn and observed ThT signal at baseline and a reduced number of fibrils by EM, further highlighting potential disruptions when labeling monomers prior to fibril formation (Figure S9). While the wild-type protein assembled primarily into a rod-like fibril polymorph, fibrils of Tet2-Et α-syn assembled into varied polymorphs (Figure 1B). Given the ready formation and abundance of unlabeled K60-Tet2-Et-α-syn fibrils, we set out to further investigate their molecular properties. K60-Tet2-Et-α-syn fibrils assembled in various buffers were generally twisting and abundant (Figure S10). Labeling of Tet2-Et-α-syn fibrils with sTCO-JF549 at a sub-stoichiometric ratio of dye to protein (Figure 1C) in their native assembly buffer yielded fluorescent fibrils whose absorbance and emission spectra matched those of free dye (Figure S3). SDS-PAGE of urea denatured fibrils (Figure S6B) showed a fluorescent band at the expected monomer molecular weight of 14 kilodaltons. A control reaction for wild-type α-syn with sTCO-JF549 showed no fluorescent band on the gel as expected due to its lack of the Tet2-Et. We also note the presence of additional fluorescent lower molecular weight bands, particularly for K96 and Y136, that may have arisen from degradation products of monomer prior to fibrillization or represent a stable fibril fragment resistant to urea and SDS denaturation. nsEM indicated the overall fibril morphologies were minimally perturbed following labeling (Figure S6C). LC-MS of K60^FL^-α-syn monomer in 10-fold molar excess of dye revealed two peaks (Figure 1D). Deconvolution of the first peak showed a mass shift of 208 Daltons corresponding to conjugation to the sTCO moiety alone (Figure 1E), while the second peak showed the expected shift corresponding to a single coupled sTCO-JF549 (Figure 1F): 645 Daltons. This corresponded to the covalent addition of one sTCO-JF549 molecule per unlabeled K60-α-syn monomer (Figure S5B). Further LC-MS analysis of all four nCAA α-syn monomers at a 1:1 molar ratio of dye to protein showed conjugation efficiency of full-length protein in the range of 10-15% under our reaction conditions (Figure S11).

**Figure 1.**
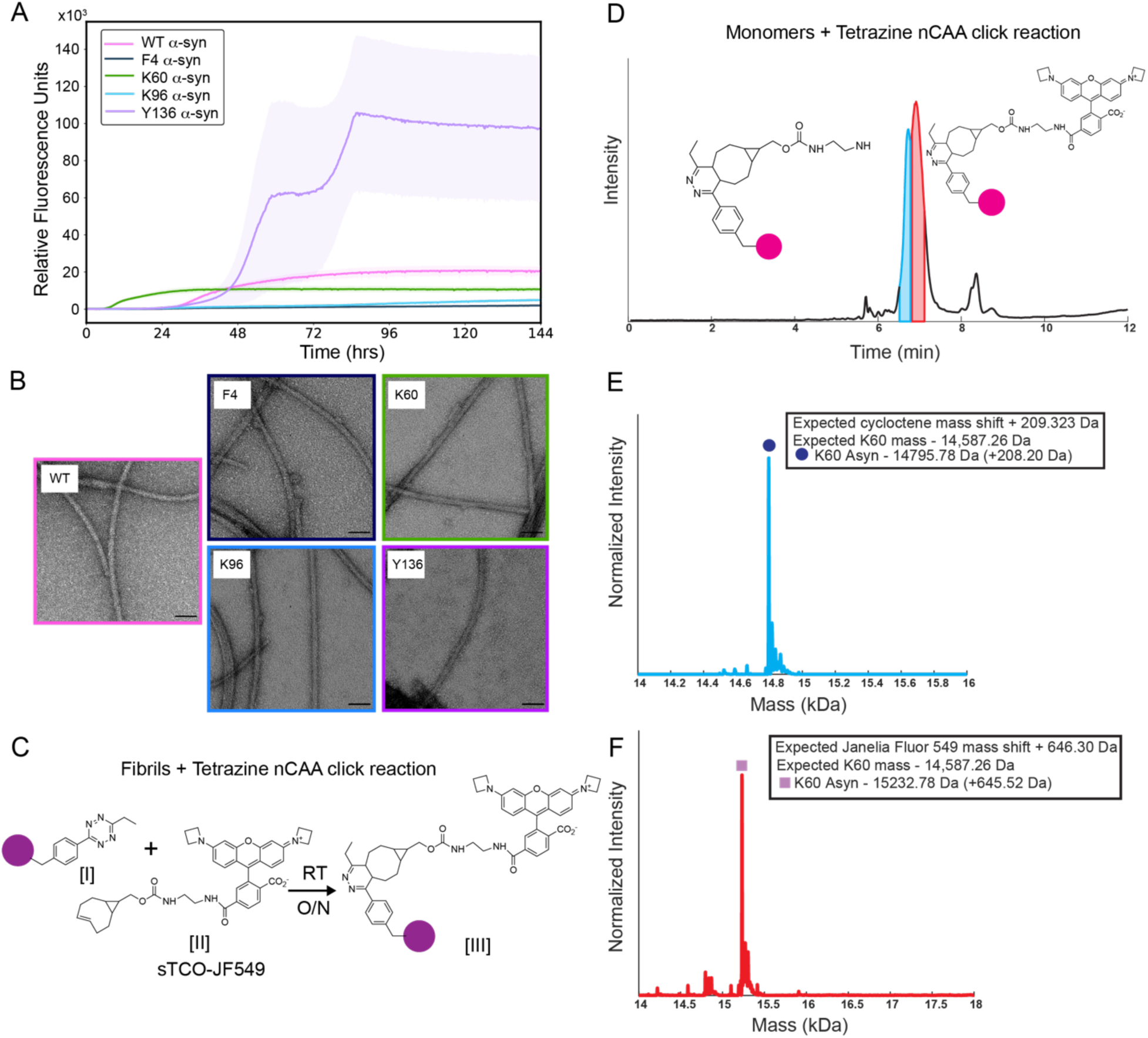
nCAA-mediated fluorophore labeling of α-syn fibrils. **(A)** Thioflavin T fibril aggregation kinetics of wild-type α-syn and that with Tet2-Et incorporated at F4, K60, K96, Y136. **(B**) Negative stain EM images of fibrils at the endpoint of the Thioflavin T assay in (A). Scale bars: 50 nm **(C)** Schematic of Tet2-Et-α-syn [I] fibril labeling with sTCO-JF549 [II] to form the fibril conjugate [III]. **(D)** LC-MS of K60^FL^-α-syn monomer. TIC elution peaks used for deconvolution are highlighted in blue and red, corresponding to the strained cyclooctene moiety alone (lacking the fluorescent group), and the full conjugation of JF549, respectively. Integrated deconvolution peaks indicating a mass corresponding to monomer conjugated to only the sTCO moiety (blue) **(E)** or the full sTCO-JF549 (red) **(F)**.

### cryoEM structure of JF549-Tet2-Et-α-syn fibrils

To better understand the properties of K60 JF549-Tet2-Et-α-syn (K60^FL^-α-syn) fibrils, we determined their structure by cryoEM. Fibrils suspended in assembly buffer were vitrified and imaged on cryoEM grids. Selection and sub-classification of fibril segments from 7815 images ultimately yielded ∼90,000 unique particles whose class averages showed striated features corresponding to ∼4.8Å spacings along the length of the fibril (Figure 2A). Those particles were used to generate an *ab initio* model of the fibril that was subsequently refined to yield a single protofilament (C1) map with a helical twist of −1.08 and rise of 4.83 Å, resolved to ∼3.1 Å resolution (Table 1, Figure S12). Residues 41-99 of α-syn were modeled into the density corresponding to the ordered core of the fibril and indicated observable density for the solvent-facing Tet2-Et residue at position 60, withTet2-Et residues on adjacent strands stacked ∼4.8 Å apart (Figure 2B-C). Two density islands were observed flanking the fibril core, but given the limited resolution attained at those regions, they remained unmodeled.

**Table 1.** CryoEM data collection and processing parameters and statistics for the determination of K60^FL^-α-syn fibrils and psychosine-induced wild-type α-syn fibrils.

|  | **K60<sup>FL</sup>-α-syn** | **Psychosine-α-syn** |
| --- | --- | --- |
| **Data collection** |  |  |
| EM equipment | Titan Krios G4 | Titan Krios G4 |
| Voltage (KV) | 300 | 300 |
| Detector | Falcon 4i | Falcon 4i |
| Nominal Magnification | x105,000 | x130,000 |
| Pixel size (Å) | 1.2 | 0.97 |
| Total electron dose (e<sup>-</sup>/Å<sup>2</sup>) | 40 | 40 |
| Defocus range (μm) | -1.5 to -2.5 | -1.5 to -2.5 |
| **Reconstruction** |  |  |
| No. of micrographs collected | 7815 | 1,641 |
| No. of micrographs used | 5076 | 1,290 |
| Box size (pixel) | 256 | 360 |
| Inter-box distance (Å) | 77 | 87 |
| No. of segments extracted | 112,620 | 116,649 |
| No. of segments for 3D auto-refine | 37,071 | 18,458 |
| Resolution (Å) | 3.07 | 2.72 |
| FSC threshold | 0.143 | 0.143 |
| Map sharpening B-factors (Å<sup>2</sup>) | -112.1 | -84.9 |
| Symmetry for final map | C1 | P2<sub>1</sub> |
| Helical twist (°) | -1.08 | 179.41 |
| Helical rise (Å) | 4.8 | 2.5 |
| **Atomic model** |  |  |
| Initial model used | 6peo | 6h6b |
| No. of non-hydrogen atoms | 2065 | 3,650 |
| No. of protein residues | 295 | 540 |
| B-factors (Å<sup>2</sup>) | 132.02 | 49.09 |
| R.m.s deviations |  |  |
| Bonds length (Å) | 0.001 | 0.003 |
| Bonds angle (°) | 0.557 | 0.611 |
| Ramachandran plot statistics (%) |  |  |
| Favored | 94.44 | 96.15 |
| Allowed | 5.56 | 3.85 |
| Outlier | 0 | 0 |
| Rotamers outliers (%) | 0 | 0 |
| C-beta deviations (%) | 0 | 0 |
| Bad bonds (%) | 0 | 0 |
| Bad angles (%) | 0 | 0 |
| MolProbity score | 1.53 | 1.88 |
| Clash score | 3.62 | 12.89 |
| PDB code | 10RO | 9Y0S |
| EMDB code | EMD-75412 | EMD-72396 |

When compared to other α-syn structures, the geometry of the K60^FL^-α-syn fibril core is most similar to that of the H50Q narrow fibril (PDB: 6PEO): a single protofilament assembly that displays an ordered core spanning residues 36-99 (Figure 2D). The K60^FL^-α-syn core has a Cα root mean squared deviation (RMSD) of less than 1Å with respect to several other structures that all adopt a type 1M fibril fold^5^. Of those, the 1M polymorph structure (PDB: 8PK2) displays a surface adsorbed beta strand that resembles an N-terminal segment resolved in juvenile-onset synucleinopathy (JOS) fibrils (PDB: 8BVQ), and fits well into a density island (U_1_) that could not be modeled in the map. That density abuts core residues 50-57, part of β2. A second density island (U_2_) is observed adjacent to residues 60-66, β3, and to the Tet2-Et density (Figure 2).

**Figure 2.**
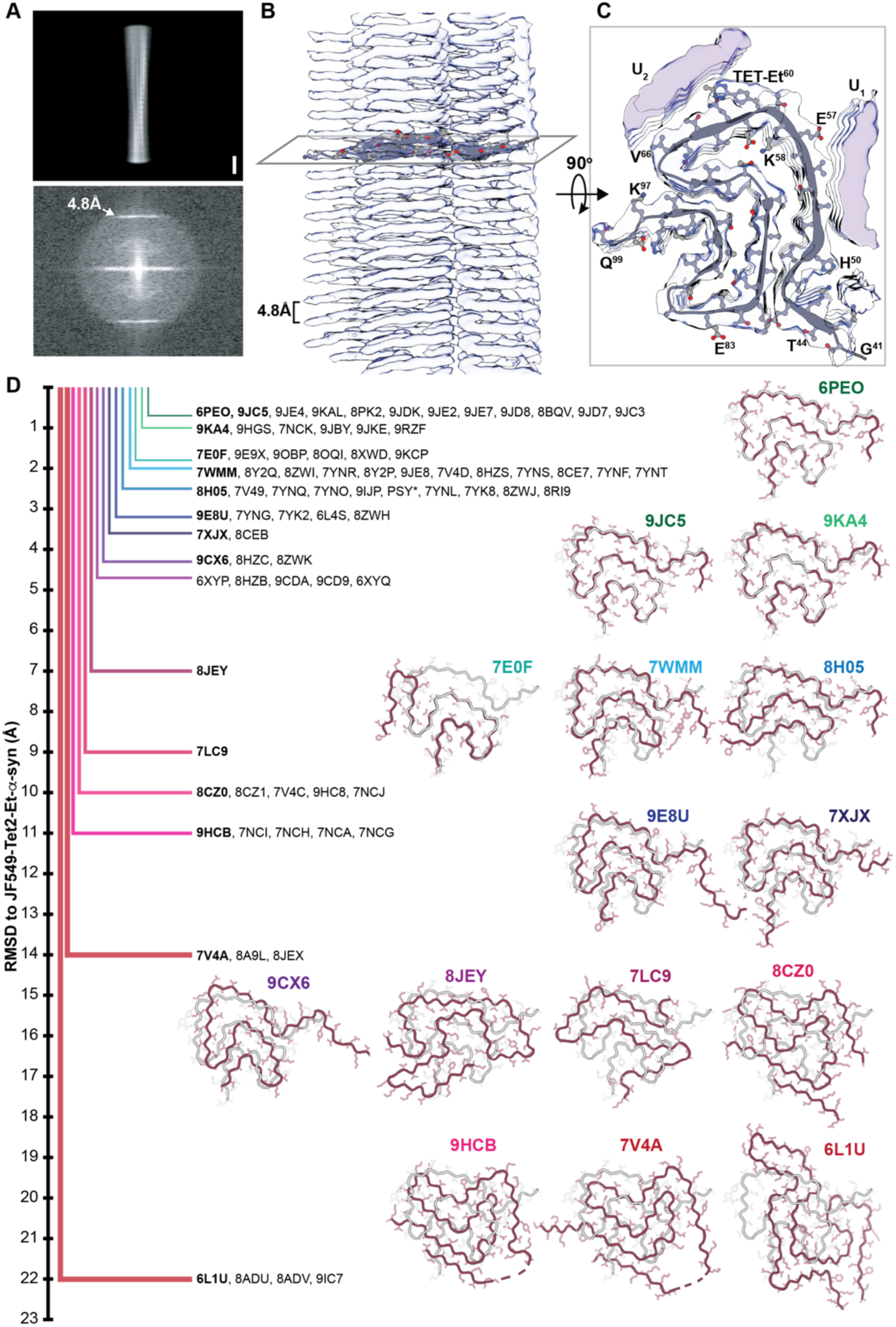
Structure of K60^FL^-α-syn fibrils. **(A)** Class average obtained from cryoEM images of K60^FL^-α-syn fibrils used for structure determination (top) and its Fourier transform (bottom); arrow points to a pronounced reflection at ∼4.8Å. Side **(B**) and top **(C)** views of the molecular model of K60^FL^-α-syn fit into its helically reconstructed, single protofilament cryoEM map. Scale bars: 50Å. **(D)** Comparison of K60^FL^-α-syn to a subset of α-syn fibril structures, listed by PDB code. Vertical axis measures RMSD of each deposited structure to K60^FL^-α-syn. Select alignments are shown with K60^FL^-α-syn as a reference (white), and query structures (burgundy) labeled by PDB code.

### Visualization of JF549-Tet2-Et-α-syn fibril-induced seeding

We evaluated the ability of K60^FL^-α-syn fibrils to seed the self-assembly of monomeric wild type α-syn and K60 Tet2-Et-α-syn with a ThT assay. We found that K60^FL^-α-syn fibrils and K60-α-syn fibrils had a reduced seeding capacity for monomeric wild type α-syn compared to wild type labeled and unlabeled fibrils (Figure 3A). Monomeric K60-α-syn alone showed accelerated fibrillization and was seeded by both K60-α-syn fibrils and K60^FL^-α-syn fibrils (Figure 3A). Wild type fibrils seeded by K60^FL^-α-syn fibrils appeared to have rod-like morphologies, while wild type fibrils seeded by K60-α-syn fibrils appeared more twisted in nsEM images (Figure 3B).

**Figure 3.**
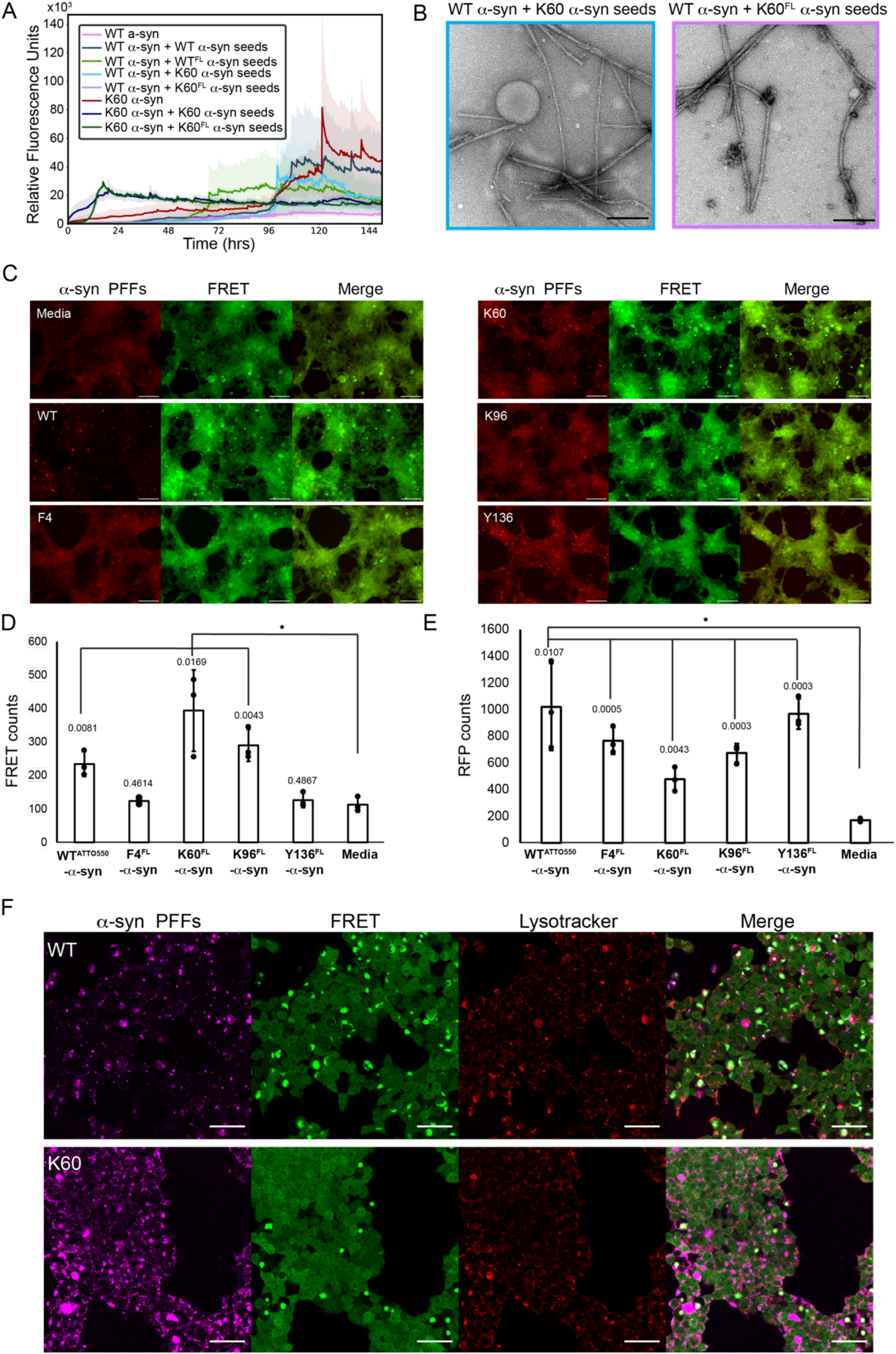
Seeding by α-syn nCAA fibril conjugates. **(A)** Thioflavin T fibril aggregation kinetics of wild-type and variant K60-α-syn monomer seeded with K60-α-syn fibrils and K60^FL^-α-syn fibrils. **(B**) Negative stain EM images of fibrils at the endpoint of the aggregation assay in (A). Scale bars: 200 nm **(C)** 24 hours post-transfection nascent aggregates, indicated by FRET signal (green), are observed in α-syn biosensor cells seeded with either ATTO-550 wild-type fibril conjugates or Tet2-Et^FL^-α-syn fibrils (red). Signal is displayed on a linear scale from 100 to 2000 (RFP) and 100 to 2000 (FRET). Scale bars: 100 µm. **(D)** Quantification of FRET signal in (C). Data are represented as mean total FRET counts for each condition, n = 3 wells. Statistical analysis: Student t test; *p ≤ 0.05 vs. media alone. Error bars = SD. F4 and Y136 did not show statistical significance compared to media alone. **(E)** Quantification of RFP signal in (C). Data are represented as mean total RFP counts for each condition, n = 3 wells. Statistical analysis: Student t test; *p ≤ 0.05 vs. media alone. Error bars = SD. The Pearson correlation coefficient for FRET and RFP counts was calculated to be −0.062. **(F)** 24 hours post-transfection 40x confocal imaging of nascent aggregates, indicated by FRET signal (green), in α-syn biosensor cells seeded with either ATTO-550 wild-type fibril conjugates labeled at a 0.1 molar ratio of dye to protein or Tet2-Et^FL^-α-syn fibrils labeled at an equal molar ratio of dye to protein, (magenta) and counterstained with lysotracker deep red (red). Scale bars: 50 µm.

K60^FL^-α-syn fibrils induced the formation of ordered aggregates in A53T α-syn biosensor cells, in similar fashion to ATTO-550 wild-type fibril conjugates. Signal from K60^FL^-α-syn fibrils was observable within cells despite sub-stoichiometric labeling ratios of dye to protein and permitted the localization of both exogenous fibrils and endogenous aggregates via RFP and FRET signals respectively (Figure 3C). Upon visual inspection, at the selected magnification, K60^FL^-α-syn fibril signal colocalized with FRET puncta to a degree that was not observed for ATTO-550 wild-type fibril conjugates. In addition, signal from labeled fibrils was punctate and distinct from that of labeled monomer conjugates in the α-syn biosensor cells (Figure S13A). Neither ATTO-550 wild-type or K60^FL^-α-syn monomer conjugates induced seeding (Figure S13B). Interestingly, we found that F4^FL^-α-syn and Y136^FL^-α-syn fibrils induced a low abundance of FRET-positive puncta in cells (Figure 3D), while K60^FL^-α-syn fibrils induced a similar abundance of FRET puncta as wild-type fibrils. Although these differences could be due to fibril uptake into cells, no obvious correlation between the number of FRET and RFP puncta was observed (Figure 3E). We speculate differences in Tet2 placement on the fibrils may modify interactions within the cell or impose a structural change that affects templating of the biosensor. Due to this difference in seeding propensity, we selected the K60 variant for further analysis. We assessed differences in cell localization between ATTO-550 wild-type fibril conjugates and K60^FL^-α-syn fibrils in the α-syn biosensor cells using lysotracker dyes. Both fibril types appeared as small puncta neighboring larger, diffuse FRET puncta in biosensor cells, and also tended to colocalize with larger lysosomal compartments (Figure 3F). Media only controls under the same detector configuration and magnification do not show fluorescent puncta in the ATTO-550 or EYFP channels and have a similar lysotracker distribution to cells seeded with fibril conjugates (Figure S14).

### The impact of Tet2-α-syn fibril induced seeding on biosensor cell health

Given the toxicity associated with amyloid fibrils^30^, we asked whether α-syn fibril conjugates promoted cellular dysfunction in biosensor cells. We analyzed the metabolic activity of cells transduced with ATTO-550 and JF549 conjugated fibrils, with the impact of fibril treatment measured against the toxicity of 5 µM methotrexate (MTX). No significant reduction in metabolic activity was observed after 24 hours of fibril-treatment in cells compared to the vehicle control (Figure S15A). Transduction of each of the four α-syn fibril conjugates, F4, K60, K96 and Y136, or wild-type ATTO550-conjugated α-syn fibrils into biosensor cells had minimal impact on their endocytic uptake of hTf-Alexa633, indicating fibril conjugates did not alter general clathrin-mediated endocytosis (Figure S15B).

### Structure informed bioconjugation of psychosine-induced α-syn fibrils

Bioorthogonal conjugation of K60-α-syn fibrils under mild buffer conditions was compatible with the presence of cofactors, including those bearing primary amines. For example, K60-α-syn fibrils could be grown and labeled in the presence of toxic lipid psychosine. This allowed direct assessment of the impact of psychosine on the aggregation propensity of α-syn, as compared to similar lipids, as monitored by ThT fluorescence and nsEM (Figure 4A). Psychosine and sphingosine accelerated fibrillization of wild type and K60-α-syn, although the lipids on their own did not form ThT-positive aggregates (Figure S16A). Similar results were observed for lipids associated with Gaucher’s disease, another LSD (Figure S16B). Wild type α-syn in the presence of psychosine formed unique polymorphs as visualized by nsEM. Psychosine-induced fibrils appeared wider than their cofactor-free counterparts with clear twisting features (Figure S17). Fibrils were prepared for cryoEM analysis in a TBS buffer, pH 8.0, that yielded a homogenous population of twisting species (Figure 4B).

**Figure 4.**
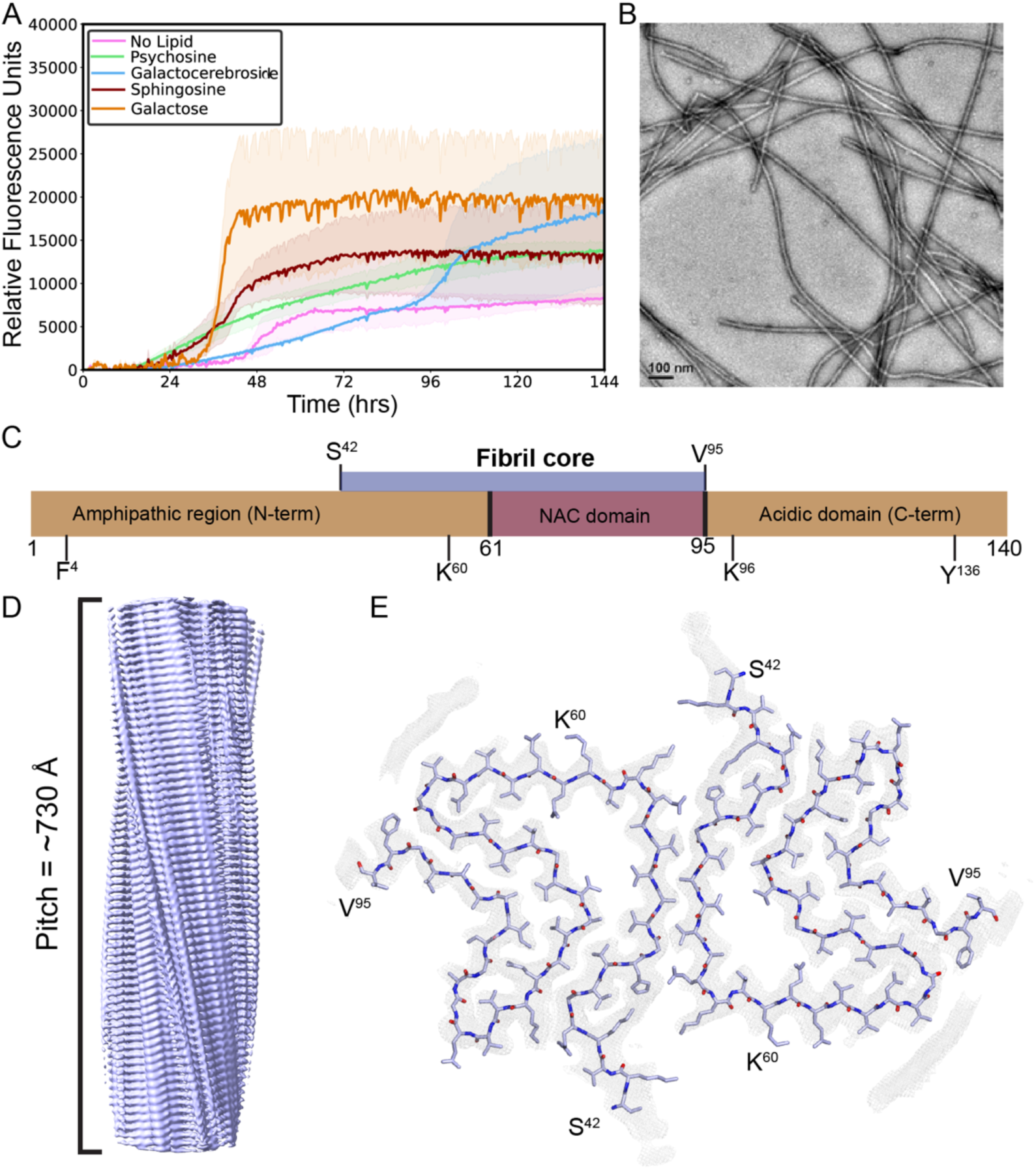
CryoEM structure of α-syn fibrils grown in the presence of psychosine. **(A)** Aggregation kinetics of wild-type α-syn fibrils self-assembled in the presence of various lipids at 10 µM. **(B)** Negative-stain EM image of wild-type α-syn fibrils grown in the presence of psychosine. Scale bar: 100 nm. **(C)** Schematic of full-length α-syn indicating the resolved fibril core, spanning residues 42-95 and the locations of sites of nCAA incorporation. **(D)** Surface view of the fibril core indicating its 730 Å pitch. **(E)** Cross-section of the fibril core with polypeptide model shown in light blue fit into the cryoEM map (gray mesh).

To determine the cryoEM structure of psychosine induced α-syn fibrils, they were diluted in buffer and centrifuged, removing larger fibril clumps. This yielded an even ice distribution across the grid with an ideal concentration of fibrils in the grid holes (Figure S18). Across 1641 micrographs (Table 1), one major fibril species could be identified and selected for 2D classification (Figure S19A). Analysis of Fourier transforms of the 2D classes showed a partial reflection at ∼2.47 Å, indicating the presence of screw-like symmetry within the ordered fibril core (Figure S19B). 2D classes of a sufficiently large box size to span a full crossover were used to generate initial symmetry parameters and a map from 3D refinement (Figure S19C). Through multiple rounds of 3D classification and 3D refinement, a map at 2.72 Å resolution was obtained that showed two protofilaments related by pseudo 2_1_ symmetry with a helical twist of 179.41° and a helical rise of 2.5 Å (Figure 4D, S20). The psychosine-induced α-syn fibril core was clearly resolved, spanning residues S42 to V95 (Figure 4C), which adopted a Greek key topology similar to type 1A fibrils, and, in particular, the rod fibril polymorph (PDB: 6CU7). An unresolved island of density located near the C-terminus was too poorly resolved for model building (Figure 4E). However, by comparison to other recombinant α-syn fibrils, it aligned well to a structure (PDB: 9C5R) that contained a similar density (Figure S21). In that structure the density was identified as an extension of the C-terminus, residues ^105^EGAPQEGILED^1^^15^ ^25^; that sequence could be partially modeled into the psychosine-induced fibril map. No other residual density was noted that could accommodate a bound psychosine molecule.

A comparison of K60^FL^-α-syn fibrils to psychosine-induced fibrils indicates a high degree of similarity between their core folds, despite their different number of protofilaments and helical symmetries (Figure S22). Both fibrils had type 1 core structures^5^, M and A for K60^FL^ and psychosine-induced, respectively, and both had satellite densities likely corresponding to ordered terminal segments (Figure S22A-C). In addition, residue 60 in the psychosine-induced fibril structure remained solvent exposed, making it compatible with conjugation in a way that would minimally impact fibril structure, as evidenced by its positioning in K60^FL^-α-syn fibrils (Figure 2, S22).

### Seeding of A53T α-syn biosensor cells with psychosine-induced JF549-Tet2-Et-α-syn fibrils

Psychosine-induced K60-α-syn fibrils exhibited the same polymorphic features as psychosine-induced wild type α-syn fibrils, suggesting K60-α-syn fibrils may have retained the psychosine induced fibril fold (Figure 5A). Psychosine K60-α-syn fibrils also readily reacted with sTCO-JF549 and could be traced when transduced into α-syn biosensor cells alongside newly seeded, FRET-positive puncta. In contrast to lipid-free fibrils, which typically appeared as pinpoint-like fluorescent puncta in biosensor cells, psychosine-induced K60^FL^-α-syn fibrils formed wider plaque-like fluorescent puncta (Figure 5B). This fibril distribution, potentially caused by the high degree of lipid-induced fibril clumping, also coincided with a similar amount of puncta formation in biosensor cells compared to its lipid free counterpart (Figure 5C). We also note that psychosine-induced fibril puncta, specifically the plaque-like morphologies, colocalized less with FRET puncta in comparison to the fluorescent puncta from their lipid-free counterpart that had a high degree of colocalization with FRET puncta (Figure 5B).

**Figure 5.**
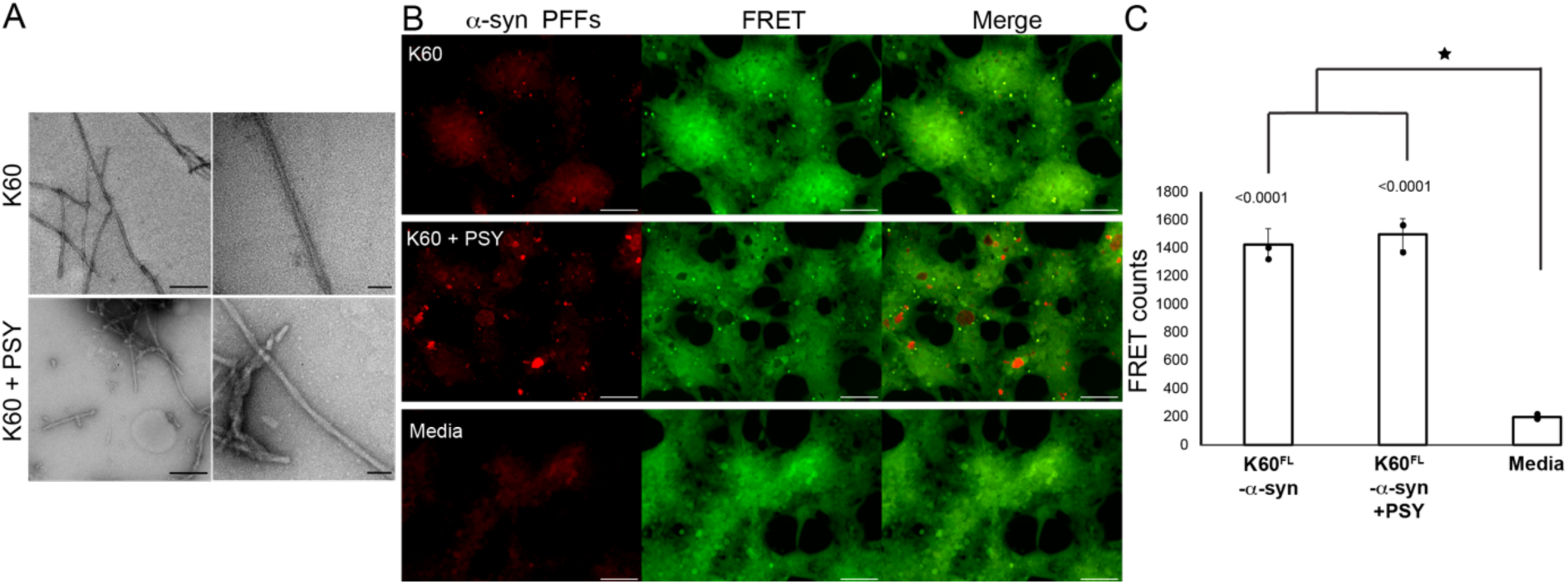
K60^FL^-α-syn fibrils grown in the presence of psychosine seed α-syn biosensor cells. **(A)** Negative-stain EM images of K60^FL^-α-syn fibrils grown with and without 10 µM psychosine (PSY). Scale bars: 200 nm (left) and 50 nm (right). **(B)** 24 hours post-transfection nascent aggregates, indicated by FRET signal (green), are observed in α-syn biosensor cells seeded with K60^FL^-α-syn fibrils (red) grown with or without 10 µM psychosine (PSY). Signal is displayed on a linear scale from 50 to 200 (RFP) and 100 to 2000 (FRET). Scale bars: 100 µm. **(C)** Mean total FRET counts for α-syn biosensor cells seeded with K60^FL^-α-syn fibrils grown with or without 10 µM psychosine, n = 3 wells. Statistical analysis: Student t test; *p ≤ 0.0001 vs. media alone. Error bars = SD.

## Discussion

Bioorthogonal labeling of amyloid fibrils mediated by non-canonical amino acid incorporation enables their routine and minimally invasive surveillance. Although many avenues have been pursued for fluorescence-based tracking of amyloid aggregates in cells, including fibril-binding dyes^31^, fluorescence protein tags^32^, and covalently coupled dyes^33^, among others, the use of bioorthogonal chemistry to couple dyes onto pre-formed fibrils offers unique advantages while allowing the study of amyloid aggregation, degradation, and spatial distribution of aggregates in cells. Site-selective conjugation of small molecule fluorophores to fibrils is less sterically hindering than genetic tagging with large proteins such as GFP that may interfere with fibril templating and alter surface properties, potentially changing how aggregates interact with other proteins^34^. As now demonstrated, a biorthogonal conjugation strategy that facilitates the rapid and efficient installation of a strained cyclooctene-containing JF549 to pre-formed α-syn fibrils bearing a Tet2 nCAA allows for effective fluorescence labeling of pre-formed fibrils. The approach is compatible with solutions containing biochemical cofactors, including those bearing primary amines. That was illustrated by our ability to grow psychosine-induced α-syn fibrils, determine their structure, and track their JF549-labeled counterparts in biosensor cells.

The structure of psychosine-induced α-syn fibrils shares many similarities with previously determined recombinant α-syn fibril structures, most notably the ordered fuzzy coat density near the C-terminus of the fibril core^35^. That feature is unique to a fibril core that showed ordering of both the N and C terminal fuzzy coats of the fibril, and the fibrils themselves were reported to have unique properties, co-aggregating with tau in primary mouse neurons^25^. It is interesting to note that the structure of psychosine induced α-syn fibrils showed no density for a bound lipid. This may in fact be due to the sub-stoichiometric concentrations of lipid used during fibril growth, which were below the critical micellar concentration for psychosine^36^. An alternative explanation could be due to weak non-covalent binding of the lipid to formed fibrils, or its preferred binding to pre-fibrillar oligomers^14,37^. While psychosine-induced K60-α-syn fibrils could be bioorthogonally conjugated without an apparent change to their morphology, the presence of Tet2 at position 60 may itself alter fibril core structure, thus affecting overall fibril properties. Those effects should be mitigated by the fact that the Tet2 site in the psychosine induced fibrils would be solvent facing, as evidenced by its efficient coupling to JF549. However, a more thorough assessment of site-selective coupling could potentially improve structural fidelity and coupling efficiency, and retain wild-type fibril interactions.

Notably, the genetic code expansion and fluorophore labeling approach can be readily applied to other amyloidogenic proteins and would allow for bioconjugation *in situ*. We also envision this strategy could be applied to disease relevant folds achieved *in vitro* for α-syn or other amyloids, such as tau. This would facilitate the use of Tet2-containing amyloid fibrils for correlative light and electron microscopy, to gain both functional and structural insights into their growth, persistence, and spread.

## Data Availability

Coordinates and cryoEM density for the structure of K60^FL^-α-syn fibrils are available via PDB entry 10RO; the corresponding raw data are available via EMPIAR deposition EMPIAR-13266. Coordinates and data for α-syn fibrils grown in the presence of psychosine are available via PDB entry 9Y0S and the corresponding raw data available via EMPIAR deposition EMPIAR-12965.

## Acknowledgments

We thank Dr. Duilio Cascio (UCLA) for technical assistance. The work was supported by NIH-NIGMS Grant R35GM128867 to J.A.R., R35GM145286 to J.A.L., the Ruth L. Kirschstein National Research Service Award program (GM007185) for C.W.F., T32GM145388, with equipment purchased with award 1S10OD016387-01, and was also performed as part of STROBE, an NSF Science and Technology Center through Grant DMR-1548924. J.A.R. was supported as a Packard Fellow. For LC-MS measurements, we acknowledge Yu Chen and UCLA’s Molecular Instrumentation Center (MIC). Instrument access was purchased from the Electron Imaging Center for Nanosystems (EICN) at the University of California, Los Angeles’s California for NanoSystems Institute (CNSI) (RRID:SCR_022900) (NIH S10OD032459 to ZHZ). Confocal laser scanning microscopy was performed at the CNSI Advanced Light Microscopy/Spectroscopy (ALMS) Laboratory and Leica Microsystems Center of Excellence at the California NanoSystems Institute at UCLA (RRID:SCR_022789) with funding support from NIH Shared Instrumentation Grant S10OD025017 and NSF Major Research Instrumentation grant CHE-0722519. This work was facilitated by the GCE4All Biomedical Technology Optimization and Dissemination Center supported by National Institute of General Medical Science grant RM1-GM144227.

## Author Contributions

R.A.J. conceptualization, methodology, formal analysis, investigation, writing, visualization; S.W. methodology, investigation; G.F. conceptualization, methodology, resources, validation; C.W.F. methodology, investigation, writing, visualization; L.J.T. methodology, formal analysis, investigation; A.H. methodology, investigation; C.S. methodology, formal analysis, investigation; M.R.S. methodology, formal analysis, investigation, supervision; J.A.L. methodology, supervision, funding acquisition; J.A.R. conceptualization, methodology, formal analysis, investigation, resources, writing, visualization, supervision, project administration, funding acquisition.

## Declaration of Interests

J.A.R is an equity stake holder of MedStruc Inc. All other authors declare no competing financial interests.

## Materials and Methods

### Protein purification

Full-length α-syn wild-type and nCAA variants were expressed and purified according to a published protocol^38^. For wild-type α-syn, BL21 cells (DE3) are transformed with the plasmid, pRK172. For nCAA mutants, BL21 cells (DE3) are double-transformed with the following plasmids: (1) pAJE-66E7: machinery plasmid expressing the Tet2-Et aminoacyl tRNA synthetase/tRNA (RS/tRNA) pair and (2) pET28-asyn^TAG^: plasmid α-syn with TAG stop codon at specified Tet2-Et position. Overnight Express ™ Instant LB Medium autoinduction media was prepared and starter culture was added such that the starting OD = 0.05. Tet2-Et was dissolved in DMSO and added to the autoinduction media to a final concentration of 0.5mM. Cultures are shaken at 250 rpm at 37C for 20-24 hours. The collected bacteria were lysed with a probe sonicator for 10 minutes (3s on/3s off, 60% amplitude) in an iced water bath. After centrifugation, the soluble fraction was boiled at 100°C for 20 minutes and cooled on ice. After centrifugation, the soluble fraction was precipitated with ammonium sulfate and the resulting pellet was resuspended and dialyzed overnight against 20 mM Tris-HCl pH 8.0. The protein was loaded onto a Hitrap Q HP column for anion exchange chromatography and eluted using 20 mM Tris-HCl, 1 M NaCl pH 8.0. Purified protein was dialyzed against water and concentrated to a minimum concentration of 3 mg/ml and stored at 4°C. The concentration of protein was determined using a Pierce BCA protein assay kit.

### Fibrillization

Wild-type a-syn and nCAA α-syn fibrils were prepared under the same conditions: 175 µM monomer, 15 mM tetrabutylphosphonium bromide, orbital shaking at 37°C for 6 days. For fibrils grown in the presence of the lipid cofactor, psychosine (Matreya), the lipid was added to monomer prior to shaking to a final concentration below the critical micelle concentration, 5-10 µM.

### Fluorescence labeling

#### ATTO-550 labeling

Preformed wild-type α-syn fibrils were dialyzed against bicarbonate buffer pH 9.5. ATTO-550 NHS ester fluorophore was dissolved in DMSO and added to fibrils at a sub stoichiometric molar ratio. The reaction mixture was agitated at room temperature for 2 hours while being protected from light. Fibrils were dialyzed back into their original growth buffer with multiple exchanges to remove excess free dye. For wild-type α-syn monomer conjugates, the ATTO-550 was added either at a molar ratio (dye to protein=0.2) for *in vitro* and cell experiments, molar ratios (dye to protein=0.1) or (dye to protein=1) for pH dependence labeling, and at molar ratios (dye to protein=1) or (dye to protein=10) for LC-MS experiments. The reaction mixture was agitated at room temperature for 2 hours and dialyzed overnight.

#### sTCO-JF549 labeling

Strained trans-cyclooctene JF549 fluorophore was dissolved in DMSO and added to Tet2-Et containing α-syn fibrils at a sub stoichiometric molar ratio (dye to protein=0.2). The reaction mixture was agitated at room temperature overnight while being protected from light. Fibrils were again dialyzed to remove excess dye. For Tet2-Et α-syn monomer conjugates, the sTCO-JF549 was added either at a molar ratio (dye to protein=0.2) for *in vitro* and cell experiments or at molar ratios (dye to protein=1) or (dye to protein=10) for LC-MS experiments unless specified otherwise. The reaction mixture was agitated at room temperature for 2 hours and dialyzed overnight.

Fibrils were analyzed for labeling using 4-12% Bis-Tris gel in 1x MES buffer. Gels were imaged using an Azure imager in the Cy3 channel.

### Absorbance/Emission spectra

Absorbance and Emission spectra for fibril and monomer conjugates were acquired on a Varioskan Lux. Absorbance values were collected through wavelengths 300-800 nm. For emission spectra, conjugates were excited with 472 nm and emission values were collected through wavelengths 490-800 nm.

### Electron microscopy imaging

Negative stain electron microscopy grids are made using F/C 300 mesh grids and stained using 2% uranyl acetate. Images are taken using a Tecnai T12 transmission electron microscope (T12) at 18500 X magnification.

### Thioflavin T assays

Purified α-syn monomers 175 μM were mixed with 20 μM ThT and added into a 96-well-plate. Samples were incubated at 37 °C for 6 days with 600 r.p.m. orbital shaking. The ThT signal was monitored using the FLUOstar Omega Microplate Reader (BMG Labtech) at an excitation wavelength of 440 nm and an emission wavelength of 490 nm.

### α-syn biosensor cell assays

Based on a published protocol^10^ α-syn biosensor cells, HEK293T cells expressing A53T a-syn tagged with CFP and YFP were cultured in DMEM supplemented with 10% FBS, 1% penicillin/strep, and 1% Glutamax. Cell cultures were maintained in a humidified atmosphere at 5% CO2 and 37°C. Cells were plated in 96 well plate at a density 20,000-30,000 cells per well for adherent cell growth. Cells were incubated at 5% CO2 and 37°C until a confluency of 70% was reached. Fibrils were sonicated for 5 minutes and diluted into a mixture of Optimem/Lipofectamine 3000 and incubated for 30 minutes at room temperature before adding to wells. Cells were incubated with transduction complexes for either 24 or 48 hours before imaging.

### Seeding quantification

To quantify seeding, FRET fluorescence events corresponding to single cells in stitched Evos images were counted using <u>MATLAB</u>. To quantify internalized fibrils, RFP fluorescence events associated with cells in stitched Evos images were counted using <u>MATLAB</u>. Average counts across replicate wells were computed for each condition along with their standard deviation. Student t-tests were evaluated for each experimental condition tested.

### WST-1 cell viability assay

HEK293T α-syn biosensor cells were plated in a 96 well plate at a density of 20,000 cells per well for adherent cell growth in 10% DMEM. Cells were incubated at 5% CO2 and 37°C until a confluency of 70% was reached. Fibrils were sonicated for 5 minutes and diluted into a mixture of Optimem/Lipofectamine 3000 and incubated for 30 minutes at room temperature before adding to wells. Controls performed included: no cells, Optimem/Lipofectamine 3000 (OM) vehicle, methotrexate (MTX) [5 µM] and DMSO vehicle (0.23% concentration in control wells). 24 hours post incubation, 5 µL WST-1 was added to wells and incubated at 37°C for 2 hours. Absorbance was measured at 440 nm for readout on metabolic activity.

### CryoEM collection and processing

#### cryoEM analysis of JF549-Tet2-Et-α-syn fibrils

sTCO-JF549-labeled (dye:protein =0.2) K60 α-syn fibrils were prepared according to our nCAA labeling method after being grown in Tris-HCl, NaCl pH 8.0, and were used for cryoEM experiments; 1.6 µL of fibril solution was applied to each side of C flat Holey Carbon 1.2/1.3 300 mesh grids. Excess liquid was blotted in a Vitrobot Mark IV set to 100% humidity/4°C and plunge frozen in liquid ethane. CryoEM data were acquired in an energy filtered G4 Krios microscope equipped with a Falcon 4i detector, operated at 300 kV and 105,000x magnification, yielding a pixel size of 1.2 Å. Data were collected using the EPU software, targeting 3 shots per grid hole.

We performed all data processing using Relion 4.0. Images collected in EER format were motion corrected to yield single frame MRC files from which fibrils were manually picked to create a training set for automated picking through all images in crYOLO^39^. 2D classification performed with segments extracted with a large (864 pixel) box size revealed two major species: one straight non-twisting and one twisting. Segments were re-extracted with a small (360 pixel) box size. These particles were high pass filtered (200 Å) and good 2D classes that showed 4.8 spacing were chosen for further refinement. We performed 3D classification with a cylinder model as the starting reference with K=4 and manual control of the tau regularization parameter and healpix order with helical symmetry searches to reach a resolution of ∼4 −5 Å. We re-extracted with a 256 pixel box and used the optimized helical parameters from previous 3D classification in subsequent 3D classification rounds to select for particles that contributed to the highest resolution class. This class was used as the initial model for 3D refinement (low pass filtered 30 A) on the final subset of particles to obtain the final reconstruction. The resulting model underwent CTF refinement to further improve the map. A final refined helical twist of 1.08° was derived from the auto-refine map and a helical rise of 4.8 Å. The final resolution was ultimately calculated to be 3.07 Å from gold-standard Fourier shell correlations at 0.143 between two independently refined half-maps.

### cryoEM analysis of psychosine-induced α-syn fibrils

α-syn fibrils grown with 5 µM psychosine in Tris-HCl, NaCl pH 8.0 were used for cryoEM analysis; 1.6 µL of fibril solution was applied to each side of Quantifoil 1.2/1.3 300 mesh grids. Excess liquid was blotted in a Vitrobot Mark IV set to 100% humidity/4°C and plunge frozen in liquid ethane. CryoEM data were acquired in an energy filtered G4 Krios microscope equipped with a Falcon 4i detector, operated at 300 kV and 130,000x magnification, yielding a pixel size of 0.97 Å. Data were collected using the EPU software, targeting 3 shots per grid hole.

We performed all data processing using Relion 4.0. Images collected in EER format were motion corrected to yield single frame MRC files, high pass filtered (200 Å), from which fibrils were manually picked and subjected to rounds of 2D classifications. 2D classification performed with segments extracted with a large (864 pixel) box size revealed two major species: one straight non-twisting and one twisting. Classes of the twisting polymorph were selected to generate initial models of different crossovers with relion_helix_inimodel2d. Segments were re-extracted with a small (360 pixel) box size. Rounds of 3D refinement were run with these segments and the initial models re-scaled to select the best initial crossover value. The map from the 750 Å crossover refinement was used as the starting reference for 3D classification. The twist and rise were deduced from multiple rounds of 3D classifications and refinement with helical symmetry searches. The resulting model underwent CTF refinement to further improve the map. A final refined helical twist of 179.405° was derived from the auto-refine map and a helical rise of 2.50 Å. During post-processing a corrected pixel size of 0.93 Å was inputted based on a 2.4 Å rise from the P2_1_ symmetry. The final resolution was ultimately calculated to be 2.72 Å from gold-standard Fourier shell correlations at 0.143 between two independently refined half-maps.

### Model building

#### Modeling of JF549-Tet2-Et-α-syn fibrils

In ChimeraX coordinates from PDB: 6peo was coarsely fit into the post-processed map. The model was further refined in Coot through manual refinement to accurately fit the model into the map. An ethyl-substituted 1,2,4,5 tetrazine nCAA (Tet2-Et) monomer was added to the structure by modifying coordinates of a methyl-substituted 1,2,4,5 tetrazine nCAA (Tet2-Me) monomer from PDB: 6NJ2 with Mercury. These coordinates were run through phenix.elbow to generate geometry restraints. The monomer was refined with real space refinement to fit the extended density replacing lysine 60. Symmetry operations were applied to gain 5 copies of the protofilament. Further refinement was done in PHENIX and through rounds of adjustments in Coot and real space refinement in PHENIX a final model was achieved that resulted in no rotamer outliers and optimal Ramachandran angles.

#### Modeling of psychosine-induced α-syn fibrils

In ChimeraX coordinates from PDB: 6h6b was coarsely fit into the post-processed map. The model was further refined in Coot through manual refinement to accurately fit the model into the map. Symmetry operations were applied to gain 5 copies of each protofilament for 10 copies total. Further refinement was done in PHENIX and through rounds of adjustments in Coot and real space refinement in PHENIX a final model was achieved that resulted in no rotamer outliers and optimal Ramachandran angles.

### Widefield fluorescence microscopy

Wide field fluorescence images were collected on an EVOS M7000 microscope equipped with DAPI, RFP, CFP/YFP (FRET) light cubes and a 20x objective. All images were acquired at that magnification and with that filter set.

### Confocal microscopy

A Leica TCS-SP8 Confocal Microscope was used for imaging fixed cells using a 40x dry or 63x oil immersion objective, and laser illumination at 408 (DAPI), 488 nm (EYFP), 552 nm (ATTO-550), and 638 (Alexa Fluor 647) with emission detected from 410-483 nm (DAPI), 493-605 nm (EYFP), 643-789 nm (ATTO-550), and 643-789 nm (Alexa Fluor 647).

### Mass spectrometry

For LC-MS analysis of conjugates, WT and K60 α-syn monomers were labeled with ATTO-550 and sTCO-JF549, respectively, at a 10-fold molar excess of dye to protein. Excess dye was dialyzed out overnight. The WT sample or the K60 sample was adjusted to 1 mg/mL determined by nanodrop A280. 3uL or 8uL were injected onto an PLRP-S (Agilent 100Å, 2.1 x 50 mm, 5 µm reversed-phase HPLC) column. Proteins were eluted by ACN gradient from 0% to 95% with a constant 0.1% FA over 12 minutes with a flow rate of 0.6ml/min. All LC-MS experiments were conducted on an Agilent 6530 Q-TOF LC/MS system in positive ion mode coupled with 1260 Infinity LC system. For the K60 sample, the observed mass shift observed for species consistent with the cyclooctene-only addition was +208.2 Da, and the mass shift for the full fluorescent probe was +645.5 Da, as expected.

For LC-MS analysis of F4, K60, K96 and Y136 conjugates, α-syn monomers were labeled with sTCO-JF549 in equal molar amounts of dye to protein. Samples were incubated at 4°C overnight then injected onto an PLRP-S (Agilent 100Å, 2.1 x 50 mm, 5 µm reversed-phase HPLC) column. Proteins were eluted by ACN gradient from 0% to 95% with a constant 0.1% FA over 12 minutes with a flow rate of 0.6ml/min. All LC-MS experiments were conducted on an Agilent 6530 Q-TOF LC/MS system in positive ion mode coupled with the 1260 Infinity LC system housed in the UCLA Molecular Instrumentation Center.

## Notes

### Summary of Updates

This version has been revised with additional results and an additional structure deposited under DPB 10RO.

## References

1. Spillantini, M. G., Schmidt, M. L., Lee, V. M.-Y., Trojanowski, J. Q., Jakes, R., and Goedert, M. (1997) α-Synuclein in Lewy bodies. Nature. 388, 839–840

2. Dickson, D. W., Liu, W.-K., Hardy, J., Farrer, M., Mehta, N., Uitti, R., Mark, M., Zimmerman, T., Golbe, L., Sage, J., Sima, A., D’Amato, C., Albin, R., Gilman, S., and Yen, S.-H. (1999) Widespread Alterations of α-Synuclein in Multiple System Atrophy. The American Journal of Pathology. 155, 1241–1251

3. Schweighauser, M., Shi, Y., Tarutani, A., Kametani, F., Murzin, A. G., Ghetti, B., Matsubara, T., Tomita, T., Ando, T., Hasegawa, K., Murayama, S., Yoshida, M., Hasegawa, M., Scheres, S. H. W., and Goedert, M. (2020) Structures of α-synuclein filaments from multiple system atrophy. Nature. 585, 464–469

4. Yang, Y., Shi, Y., Schweighauser, M., Zhang, X., Kotecha, A., Murzin, A. G., Garringer, H. J., Cullinane, P. W., Saito, Y., Foroud, T., Warner, T. T., Hasegawa, K., Vidal, R., Murayama, S., Revesz, T., Ghetti, B., Hasegawa, M., Lashley, T., Scheres, S. H. W., and Goedert, M. (2022) Structures of α-synuclein filaments from human brains with Lewy pathology. Nature. 610, 791–795

5. Frey, L., Ghosh, D., Qureshi, B. M., Rhyner, D., Guerrero-Ferreira, R., Pokharna, A., Kwiatkowski, W., Serdiuk, T., Picotti, P., Riek, R., and Greenwald, J. (2024) On the pH-dependence of α-synuclein amyloid polymorphism and the role of secondary nucleation in seed-based amyloid propagation. eLife. 12, RP93562

6. Todd, T. W., Islam, N. N., Cook, C. N., Caulfield, T. R., and Petrucelli, L. (2024) Cryo-EM structures of pathogenic fibrils and their impact on neurodegenerative disease research. Neuron. 112, 2269–2288

7. Henderson, M. X., Cornblath, E. J., Darwich, A., Zhang, B., Brown, H., Gathagan, R. J., Sandler, R. M., Bassett, D. S., Trojanowski, J. Q., and Lee, V. M. Y. (2019) Spread of α-synuclein pathology through the brain connectome is modulated by selective vulnerability and predicted by network analysis. Nat Neurosci. 22, 1248–1257

8. Burger, D., Kashyrina, M., Van Den Heuvel, L., De La Seiglière, H., Lewis, A. J., De Nuccio, F., Mohammed, I., Verchère, J., Feuillie, C., Berbon, M., Arotcarena, M.-L., Retailleau, A., Bezard, E., Canron, M.-H., Meissner, W. G., Loquet, A., Bousset, L., Poujol, C., Nilsson, K. P. R., Laferrière, F., Baron, T., Lofrumento, D. D., De Giorgi, F., Stahlberg, H., and Ichas, F. (2025) Synthetic α-synuclein fibrils replicate in mice causing MSA-like pathology. Nature. 648, 409–417

9. Ban, T., Hamada, D., Hasegawa, K., Naiki, H., and Goto, Y. (2003) Direct Observation of Amyloid Fibril Growth Monitored by Thioflavin T Fluorescence * 210. Journal of Biological Chemistry. 278, 16462–16465

10. Holmes, B. B., Furman, J. L., Mahan, T. E., Yamasaki, T. R., Mirbaha, H., Eades, W. C., Belaygorod, L., Cairns, N. J., Holtzman, D. M., and Diamond, M. I. (2014) Proteopathic tau seeding predicts tauopathy in vivo. Proc Natl Acad Sci U S A. 111, E4376–4385

11. Vaquer-Alicea, J., Manon, V. A., Bommareddy, V., Kunach, P., Gupta, A., Monistrol, J., Perez, V. A., Tran, H. T., Saez-Calveras, N., Du, S., Batra, S., Stoddard, D., White, C. L., Joachimiak, L. A., Shahmoradian, S. H., and Diamond, M. I. (2025) Functional classification of tauopathy strains reveals the role of protofilament core residues. Sci. Adv. 11, eadp5978

12. Platt, F. M., d’Azzo, A., Davidson, B. L., Neufeld, E. F., and Tifft, C. J. (2018) Lysosomal storage diseases. Nat Rev Dis Primers. 4, 27

13. Bradbury, A. M., Bongarzone, E. R., and Sands, M. S. (2021) Krabbe disease: New hope for an old disease. Neuroscience Letters. 752, 135841

14. Smith, B. R., Santos, M. B., Marshall, M. S., Cantuti-Castelvetri, L., Lopez-Rosas, A., Li, G., Van Breemen, R., Claycomb, K. I., Gallea, J. I., Celej, M. S., Crocker, S. J., Givogri, M. I., and Bongarzone, E. R. (2014) Neuronal inclusions of α-synuclein contribute to the pathogenesis of Krabbe disease. The Journal of Pathology. 232, 509–521

15. Hatton, C., Ghanem, S. S., Koss, D. J., Abdi, I. Y., Gibbons, E., Guerreiro, R., Bras, J., International DLB Genetics Consortium, Bras, J., Guerreiro, R., Kun-Rodrigues, C., Singleton, A., Hernandez, D., Ross, O. A., Dickson, D. W., Graff-Radford, N., Ferman, T. J., Petersen, R. C., Boeve, B. F., Heckman, M. G., Trojanowski, J. Q., Van Deerlin, V., Cairns, N. J., Morris, J. C., Stone, D. J., Eicher, J. D., Clark, L., Honig, L. S., Marder, K., Serrano, G. E., Beach, T. G., Galasko, D., Masliah, E., Hardy, J., Darwent, L., Ansorge, O., Parkkinen, L., Morgan, K., Brown, K., Braae, A., Barber, I., Troakes, C., Al-Sarraj, S., Warner, T., Lashley, T., Holton, J., Compta, Y., Revesz, T., Lees, A., Zetterberg, H., Escott-Price, V., Pickering-Brown, S., Mann, D., George-Hyslop, P. St., Rogaeva, E., George-Hyslop, P. St., Clarimon, J., Lleo, A., Morenas-Rodriguez, E., Pastor, P., Diez-Fairen, M., Aquilar, M., Compta, Y., Shepherd, C., Halliday, G. M., Tienari, P. J., Myllykangas, L., Oinas, M., Santana, I., Lesage, S., Zetterberg, H., Londos, E., Lemstra, A., Walker, L., Gelpi, E., Heywood, W., Outeiro, T. F., Attems, J., McFarland, R., Forsyth, R., El-Agnaf, O. M., and Erskine, D. (2022) Prion-like α-synuclein pathology in the brain of infants with Krabbe disease. Brain. 145, 1257–1263

16. Mujumdar, R. B., Ernst, L. A., Mujumdar, S. R., Lewis, C. J., and Waggoner, A. S. (1993) Cyanine dye labeling reagents: sulfoindocyanine succinimidyl esters. Bioconjug Chem. 4, 105–111

17. Kim, Y., Ho, S. O., Gassman, N. R., Korlann, Y., Landorf, E. V., Collart, F. R., and Weiss, S. (2008) Efficient Site-Specific Labeling of Proteins via Cysteines. Bioconjugate Chem. 19, 786–791

18. Wall, A., Wills, A. G., Forte, N., Bahou, C., Bonin, L., Nicholls, K., Ma, M. T., Chudasama, V., and Baker, J. R. (2020) One-pot thiol–amine bioconjugation to maleimides: simultaneous stabilisation and dual functionalisation. Chem. Sci. 11, 11455–11460

19. Van Moorsel, M. V. A., Urbanus, R. T., Verhoef, S., Koekman, C. A., Vink, M., Vermonden, T., Maas, C., Pasterkamp, G., and Schiffelers, R. M. (2019) A head-to-head comparison of conjugation methods for VHHs: Random maleimide-thiol coupling versus controlled click chemistry. International Journal of Pharmaceutics: X. 1, 100020

20. Karpowicz, R. J., Haney, C. M., Mihaila, T. S., Sandler, R. M., Petersson, E. J., and Lee, V. M.-Y. (2017) Selective imaging of internalized proteopathic α-synuclein seeds in primary neurons reveals mechanistic insight into transmission of synucleinopathies. Journal of Biological Chemistry. 292, 13482–13497

21. Iljina, M., Garcia, G. A., Horrocks, M. H., Tosatto, L., Choi, M. L., Ganzinger, K. A., Abramov, A. Y., Gandhi, S., Wood, N. W., Cremades, N., Dobson, C. M., Knowles, T. P. J., and Klenerman, D. (2016) Kinetic model of the aggregation of alpha-synuclein provides insights into prion-like spreading. Proc. Natl. Acad. Sci. U.S.A. 10.1073/pnas.1524128113

22. Zhang, X., Wesén, E., Kumar, R., Bernson, D., Gallud, A., Paul, A., Wittung-Stafshede, P., and Esbjörner, E. K. (2020) Correlation between Cellular Uptake and Cytotoxicity of Fragmented α-Synuclein Amyloid Fibrils Suggests Intracellular Basis for Toxicity. ACS Chem. Neurosci. 11, 233–241

23. Mučibabić, M., Apetri, M. M., Canters, G. W., and Aartsma, T. J. (2016) The effect of fluorescent labeling on α-synuclein fibril morphology. Biochimica et Biophysica Acta (BBA) - Proteins and Proteomics. 1864, 1419–1427

24. Ilievski, F., Wikström, L., Borg, A., Volkov, I. L., Brandis, G., and Johansson, M. (2026) Optimization of the genetic code expansion technology for intracellular labelling and single-molecule tracking of proteins in genomically re-coded E. coli. RSC Chem. Biol. 7, 269–285

25. Sun, C., Zhou, K., DePaola, P., Li, C., Lee, V. M. Y., Zhou, Z. H., Peng, C., and Jiang, L. (2025) Structural basis of a distinct α-synuclein strain that promotes tau inclusion in neurons. Journal of Biological Chemistry. 301, 108351

26. Sokratian, A., Zhou, Y., Tatli, M., Burbidge, K. J., Xu, E., Viverette, E., Donzelli, S., Duda, A. M., Yuan, Y., Li, H., Strader, S., Patel, N., Shiell, L., Malankhanova, T., Chen, O., Mazzulli, J. R., Perera, L., Stahlberg, H., Borgnia, M., Bartesaghi, A., Lashuel, H. A., and West, A. B. (2024) Mouse α-synuclein fibrils are structurally and functionally distinct from human fibrils associated with Lewy body diseases. Sci. Adv. 10, eadq3539

27. Eddins, A. J., Pung, A. H., Cooley, R. B., and Mehl, R. A. (2024) Tetrazine Amino Acid Encoding for Rapid and Complete Protein Bioconjugation. Bio Protoc. 14, e5048

28. Poty, S., Membreno, R., Glaser, J. M., Ragupathi, A., Scholz, W. W., Zeglis, B. M., and Lewis, J. S. (2018) The inverse electron-demand Diels–Alder reaction as a new methodology for the synthesis of^225^ Ac-labelled radioimmunoconjugates. Chem. Commun. 54, 2599–2602

29. Tuttle, M. D., Comellas, G., Nieuwkoop, A. J., Covell, D. J., Berthold, D. A., Kloepper, K. D., Courtney, J. M., Kim, J. K., Barclay, A. M., Kendall, A., Wan, W., Stubbs, G., Schwieters, C. D., Lee, V. M. Y., George, J. M., and Rienstra, C. M. (2016) Solid-state NMR structure of a pathogenic fibril of full-length human α-synuclein. Nat Struct Mol Biol. 23, 409–415

30. Matveyenka, M., Rizevsky, S., and Kurouski, D. (2022) Amyloid aggregates exert cell toxicity causing irreversible damages in the endoplasmic reticulum. Biochim Biophys Acta Mol Basis Dis. 1868, 166485

31. Spehar, K., Ding, T., Sun, Y., Kedia, N., Lu, J., Nahass, G. R., Lew, M. D., and Bieschke, J. (2018) Super-resolution Imaging of Amyloid Structures over Extended Times by Using Transient Binding of Single Thioflavin T Molecules. ChemBioChem. 19, 1944–1948

32. Bäuerlein, F. J. B., Saha, I., Mishra, A., Kalemanov, M., Martínez-Sánchez, A., Klein, R., Dudanova, I., Hipp, M. S., Hartl, F. U., Baumeister, W., and Fernández-Busnadiego, R. (2017) In Situ Architecture and Cellular Interactions of PolyQ Inclusions. Cell. 171, 179–187.e10

33. Bhuskute, K. R., Kikuchi, K., Luo, Z., and Kaur, A. (2024) Visualizing Amyloid Assembly at the Nanoscale: Insights from Super-Resolution Imaging. Analysis & Sensing. 4, e202400001

34. Shillcock, J. C., Hastings, J., Riguet, N., and Lashuel, H. A. (2022) Non-monotonic fibril surface occlusion by GFP tags from coarse-grained molecular simulations. Computational and Structural Biotechnology Journal. 20, 309–321

35. Guerrero-Ferreira, R., Kovacik, L., Ni, D., and Stahlberg, H. (2020) New insights on the structure of alpha-synuclein fibrils using cryo-electron microscopy. Current Opinion in Neurobiology. 61, 89–95

36. Zulueta Díaz, Y. D. L. M., Caby, S., Bongarzone, E. R., and Fanani, M. L. (2018) Psychosine remodels model lipid membranes at neutral pH. Biochimica et Biophysica Acta (BBA) - Biomembranes. 1860, 2515–2526

37. Taguchi, Y. V., Liu, J., Ruan, J., Pacheco, J., Zhang, X., Abbasi, J., Keutzer, J., Mistry, P. K., and Chandra, S. S. (2017) Glucosylsphingosine Promotes α-Synuclein Pathology in Mutant GBA-Associated Parkinson’s Disease. J. Neurosci. 37, 9617–9631

38. Li, B., Ge, P., Murray, K. A., Sheth, P., Zhang, M., Nair, G., Sawaya, M. R., Shin, W. S., Boyer, D. R., Ye, S., Eisenberg, D. S., Zhou, Z. H., and Jiang, L. (2018) Cryo-EM of full-length α-synuclein reveals fibril polymorphs with a common structural kernel. Nat Commun. 9, 3609

39. Wagner, T., Merino, F., Stabrin, M., Moriya, T., Antoni, C., Apelbaum, A., Hagel, P., Sitsel, O., Raisch, T., Prumbaum, D., Quentin, D., Roderer, D., Tacke, S., Siebolds, B., Schubert, E., Shaikh, T. R., Lill, P., Gatsogiannis, C., and Raunser, S. (2019) SPHIRE-crYOLO is a fast and accurate fully automated particle picker for cryo-EM. Commun Biol. 2, 218

