## Supplemental tables and figures for "Leveraging structurally-informed bioorthogonal conjugation for alpha synuclein fibril surveillance"

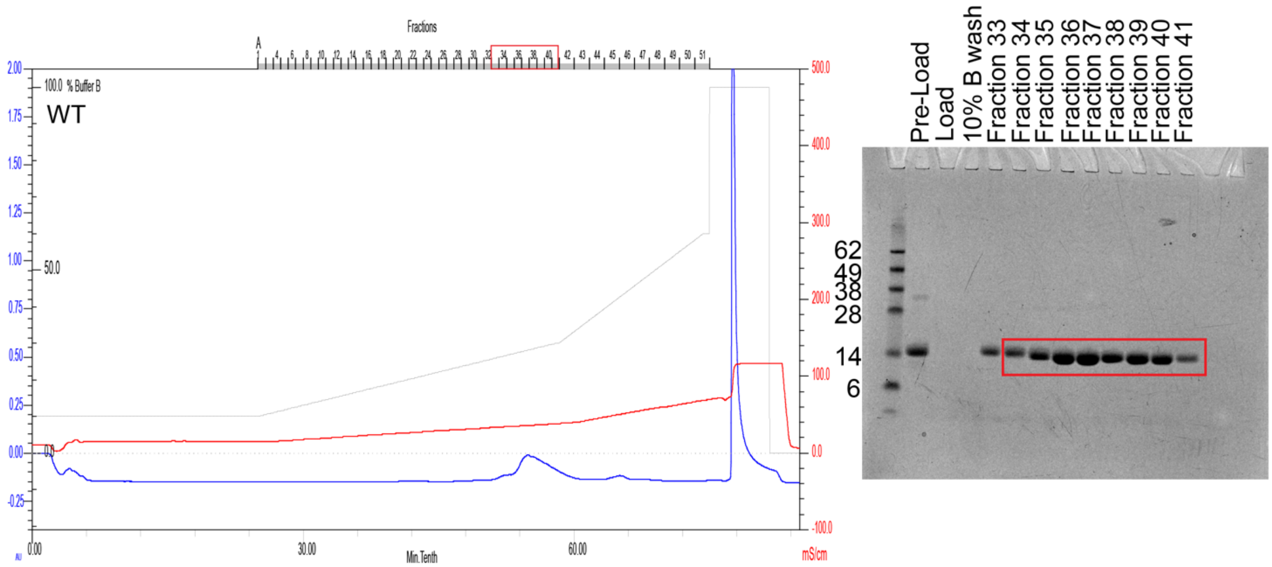


**Figure S1. Purification of recombinant wild-type α-syn monomers via anion exchange chromatography**. Chromatogram (left) showing UV peak and SDS-PAGE gel (right) showing pure fractions boxed in red that were pooled.

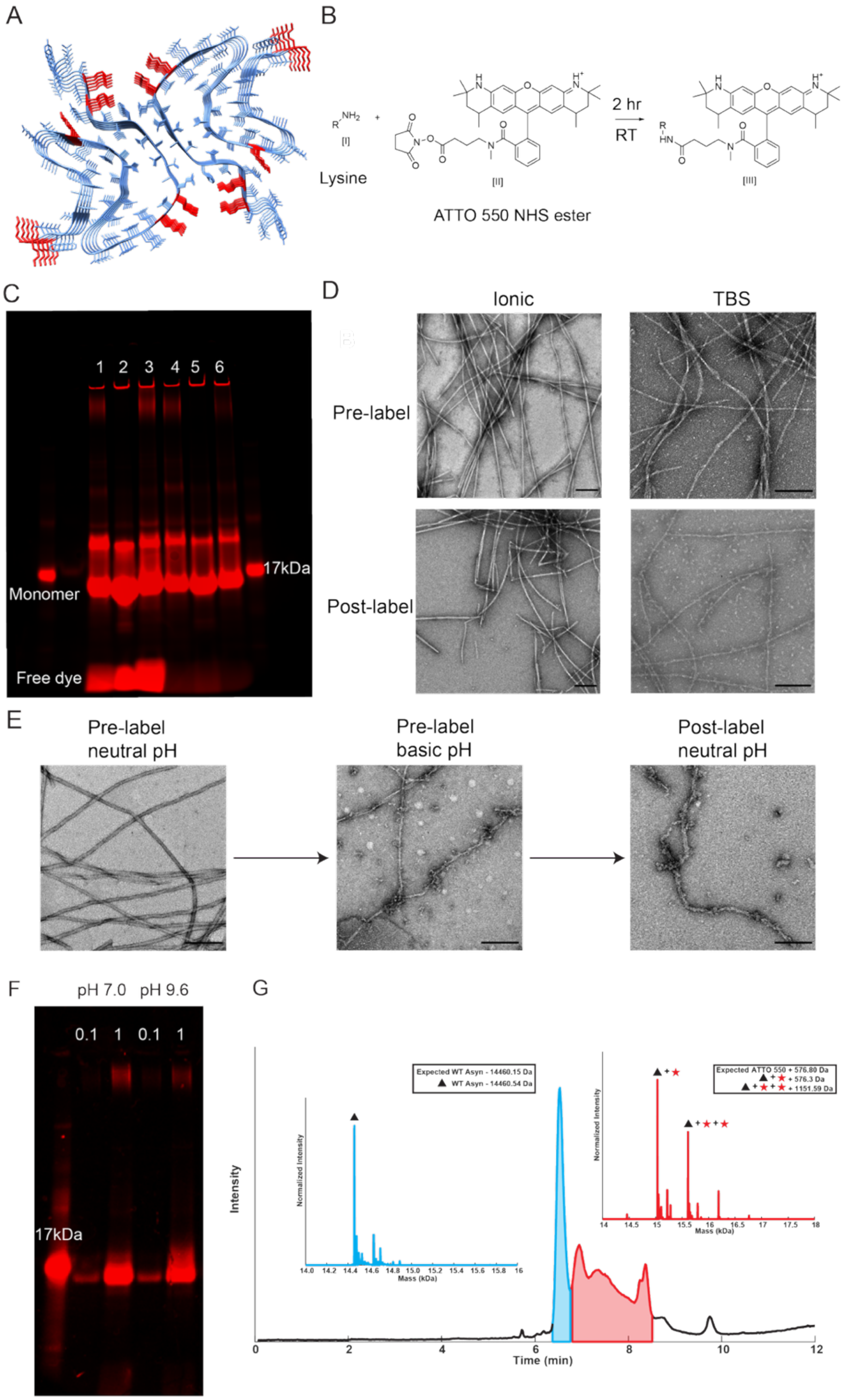


**Figure S2. Wild-type α-syn ATTO-550 labeling**. **(A)** Locations of amino acids in wild-type fibril structure (PDB: 6CU7) that have solvent exposed lysine residues. **(B)** Schematic of lysine conjugation to ATTO-550 NHS ester under basic pH conditions. **(C)** Validation of successful conjugation of fibrils with ATTO-550 via SDS-PAGE fluorescence imaging: pre-dialysis (lanes 1-3) and post-dialysis to remove free dye (lanes 4-6). **(D)** Negative-stain EM images of fibrils pre-conjugation (top row) and post-conjugation (bottom row). Scale bars are 200 nm. **(E)** ATTO-550 modified fibrils imaged by EM after 2 hour period of buffer exchange into bicarbonate buffer (pH 9.5). **(F)** Validation of α-syn ATTO-550 monomer conjugation under neutral and basic pH conditions via SDS-PAGE fluorescence imaging **(G)** LC-MS of wild-type α-syn ATTO-550 monomer conjugation. Total ion chromatogram elution peaks used for deconvolution are highlighted in blue and red, corresponding to the unconjugated protein and a series of ATTO-550 conjugated species, respectively. The blue inset shows the integrated deconvoluted mass spectrum of the unconjugated peak, revealing a mass consistent with wild-type α-syn. The red inset displays the deconvolution of the later-eluting species, indicating a distribution of conjugates corresponding to the addition of 1 or 2 ATTO-550 molecules.

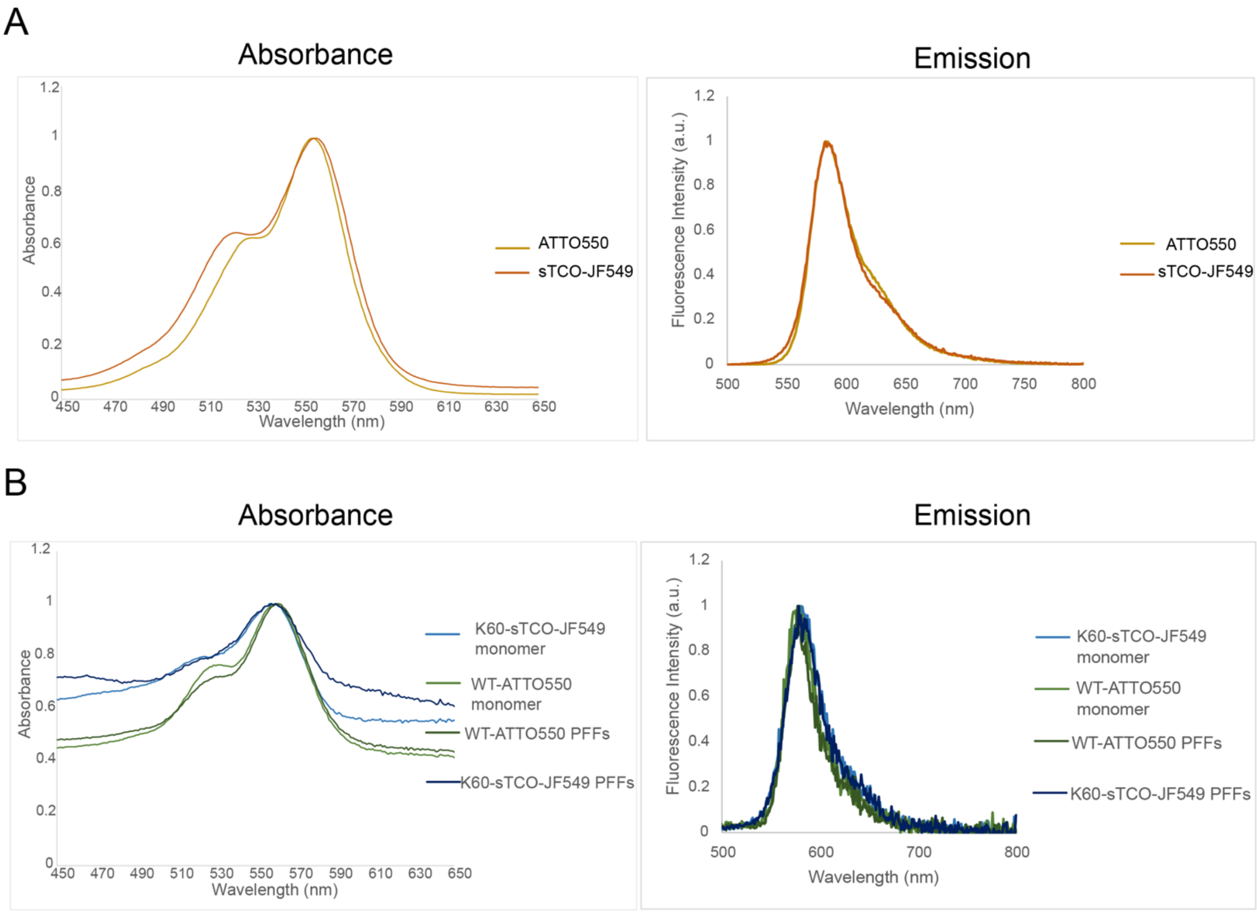


**Figure S3. Absorbance and fluorescence emission for ATTO-550 and sTCO-JF549 conjugates**. **(A)** Absorbance spectra from 450- 800 nm and emission spectra from excitation at 472 nm for free ATTO-550 (gold) and sTCO-JF549 (orange). **(B)** Absorbance spectra from 450-800 nm and emission spectra from excitation at 472 nm for) for protein conjugates: WT monomer (light green), WT fibrils (dark green), K60 monomer (light blue), and K60 fibrils (dark blue).

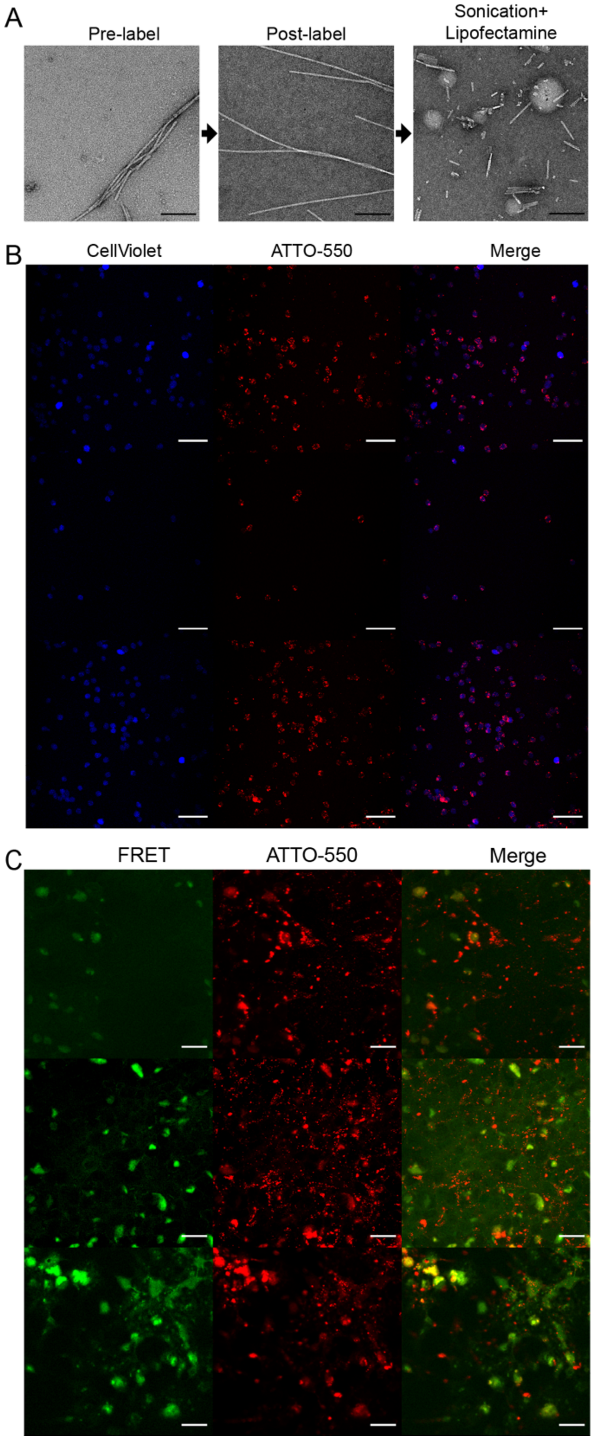


**Figure S4. Surveilling fluorophore-conjugated α-syn fibrils in HEK293T cells**. **(A)** Negative-stain EM micrographs of α-syn fibrils pre and post labeling via ATTO-550 conjugation and post-labeling sonication and lipofectamine packaging for delivery into cells. Scale bar: 200 nm. **(B)** CellTrace™ Violet (blue) labeled HEK293T cells were seeded with wild-type α-syn fibrils conjugated to ATTO-550 (red). 24 hours post-transfection cells from quadruplicate wells were pooled and fixed for confocal imaging. Three regions of interest were acquired in the DAPI and ATTO-550 channels at 20x. Scale bars: 250 nm **(C)** HEK293T α-syn biosensor cells were directly grown on coverslips prior to seeding. 48 hours post-transfection cells were fixed for confocal imaging. Three regions of interest were acquired in the ATTO-550 and EYFP channels at 63x. FRET signal (green) indicating nascent α-syn aggregates in HEK293T α-syn biosensor cells seeded with ATTO-550-conjugated wild-type α-syn fibrils (red). Scale bars: 100 nm.

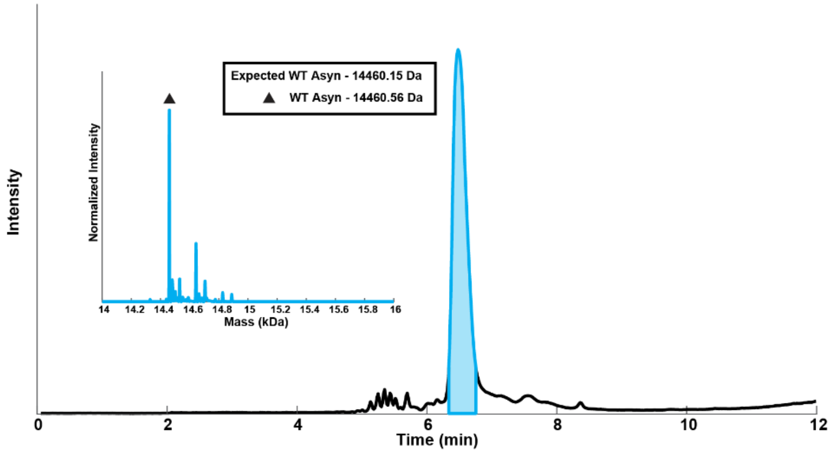


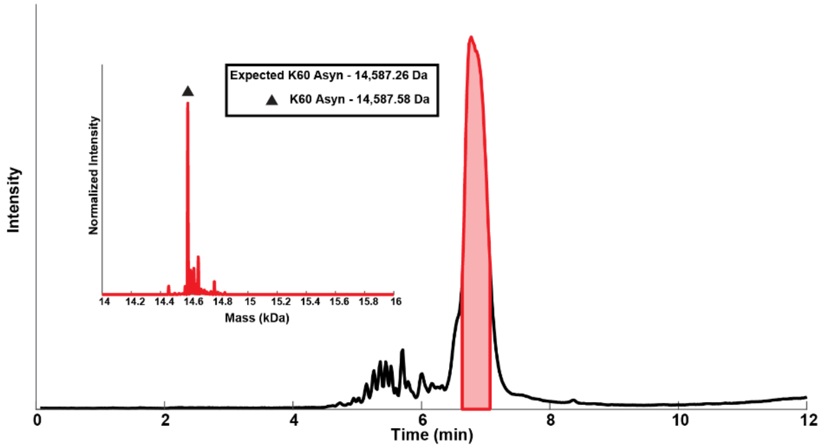


**Figure S5. LC-MS analysis of unconjugated wild-type α-syn and K60 α-syn**. The total ion chromatogram (TIC) of wild-type α-syn is shown, with the elution peak highlighted in blue indicating the time window averaged for deconvolution. The inset displays the integrated deconvolution mass spectrum from this region, revealing the observed molecular weight, indicated by the black triangle, matches with the expected dominant species of wildtype α-syn **(A)**. The TIC of K60 α-syn is shown, with the elution peak highlighted in red indicating the time window averaged for deconvolution. The inset displays the integrated deconvoluted mass spectrum from this region, revealing the observed molecular weight, indicated by the black triangle, matches with the expected dominant species of K60 α-syn **(B)**.

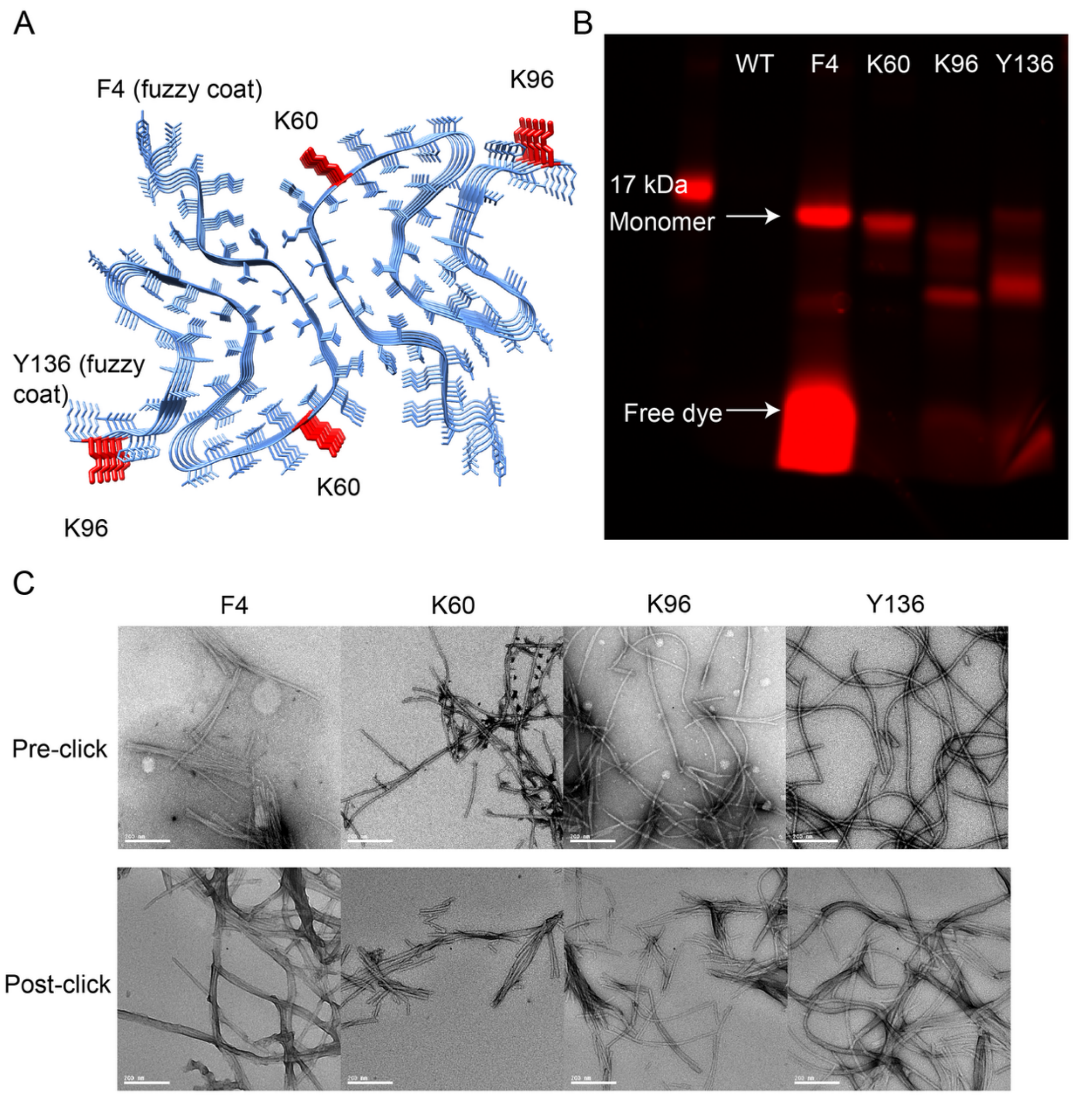


**Figure S6. Non-canonical amino acid α-syn click reaction. (A)** Locations of amino acids in wild-type fibril structure (PDB: 6CU7) that were chosen to be modified with the non-canonical amino acid, 1,2,4,5-tetrazine-ethyl (Tet2-Et). **(B)** Validation of successful conjugation of fibrils with sTCO-JF549 via SDS-PAGE fluorescence imaging. **(C)** Negative-stain EM images of fibrils pre and post labeling with click reaction. Scale bars are 200 nm.

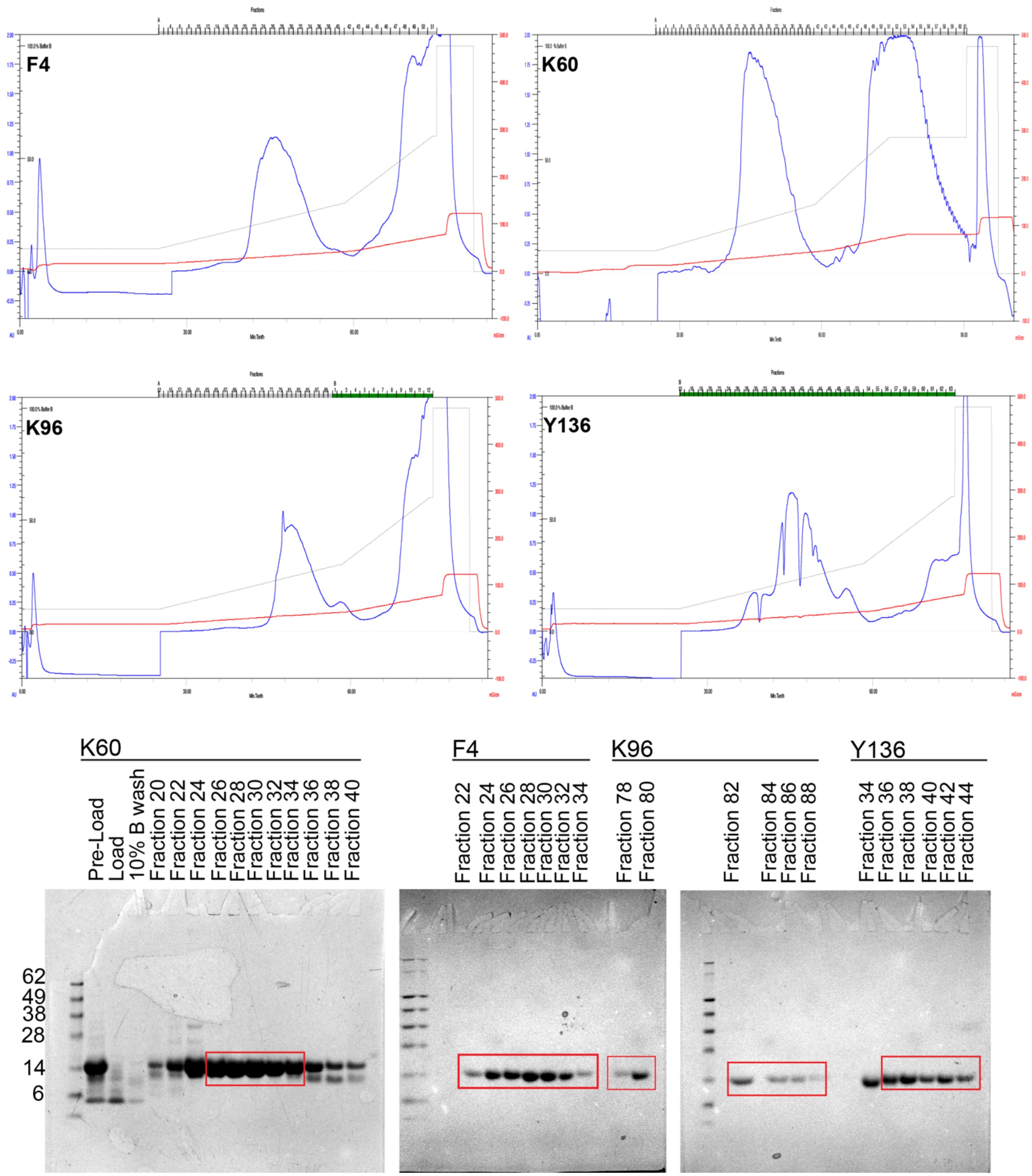


**Figure S7. Purification of recombinant Tet2-Et incorporated α-syn monomers via anion exchange chromatography**. Chromatograms showing UV peak and SDS-PAGE gels showing pure fractions boxed in red that were pooled for each Tet2-Et α-syn monomers.

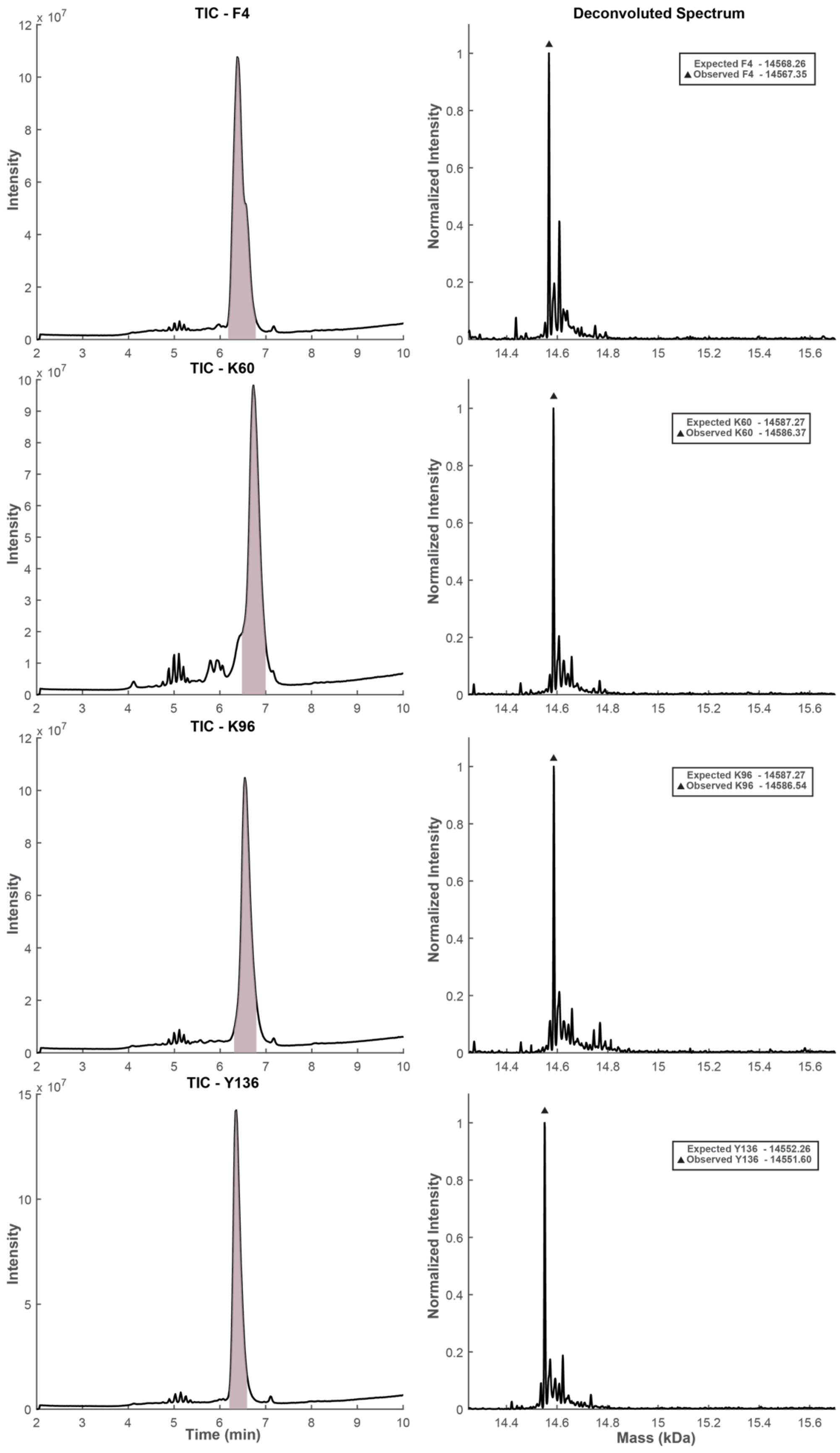

TIC - F4
Deconvoluted Spectrum
Expected F4 - 14568.26
Observed F4 - 14567.35
TIC - K60
Expected K60 - 14587.27
Observed K60 - 14586.37
TIC - K96
Expected K96 - 14587.27
Observed K96 - 14586.54
TIC - Y136
Expected Y136 - 14552.26
Observed Y136 - 14551.60
Intensity
Normalized Intensity
Time (min)
Mass (kDa)

**Figure S8. LC-MS analysis of Tet2-Et incorporated α-syn monomers.** The total ion chromatogram of each nCAA monomer with the elution peak highlighted in purple indicating the time window averaged for deconvolution (left). The integrated deconvolution mass spectrum from this region, revealing the observed molecular weights, indicated by the black triangles (right).

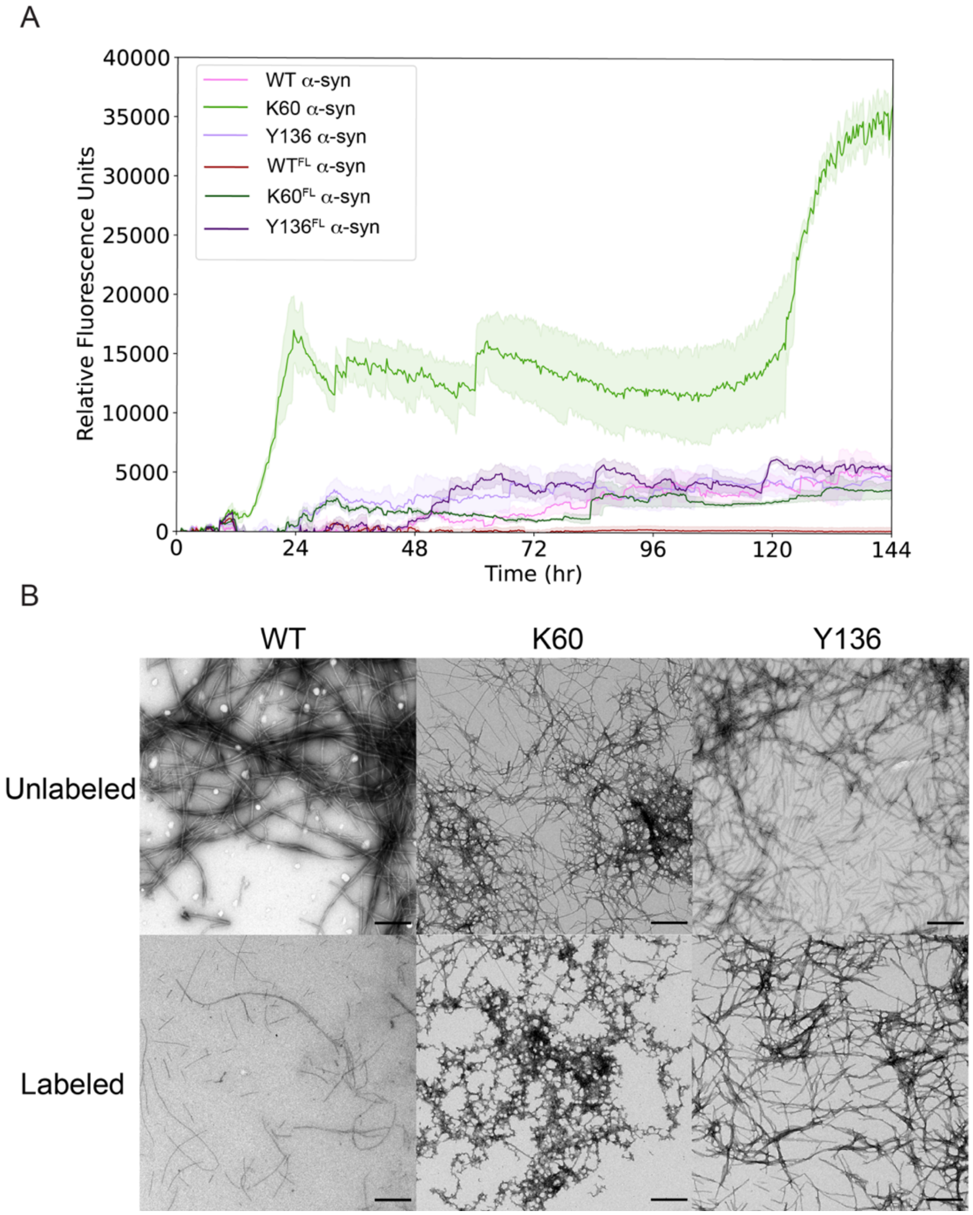


**Figure S9. Thioflavin T assay with labeled monomers and unlabeled monomers**. **(A)** Thioflavin T fibril aggregation kinetics of wild-type and variant K60 and Y136 α-syn monomers both unlabeled and labeled with their respective fluorophores, ATTO-550 for wild-type and sTCO-JF549 for variants, at a dye to protein ratio of 0.2. Fibrillization occurred in ionic additive conditions. The fibrils formed at the endpoint of the Thioflavin T assay at 144 hours were characterized by negative-stain EM **(B)** Scale bars are 0.5 μm.

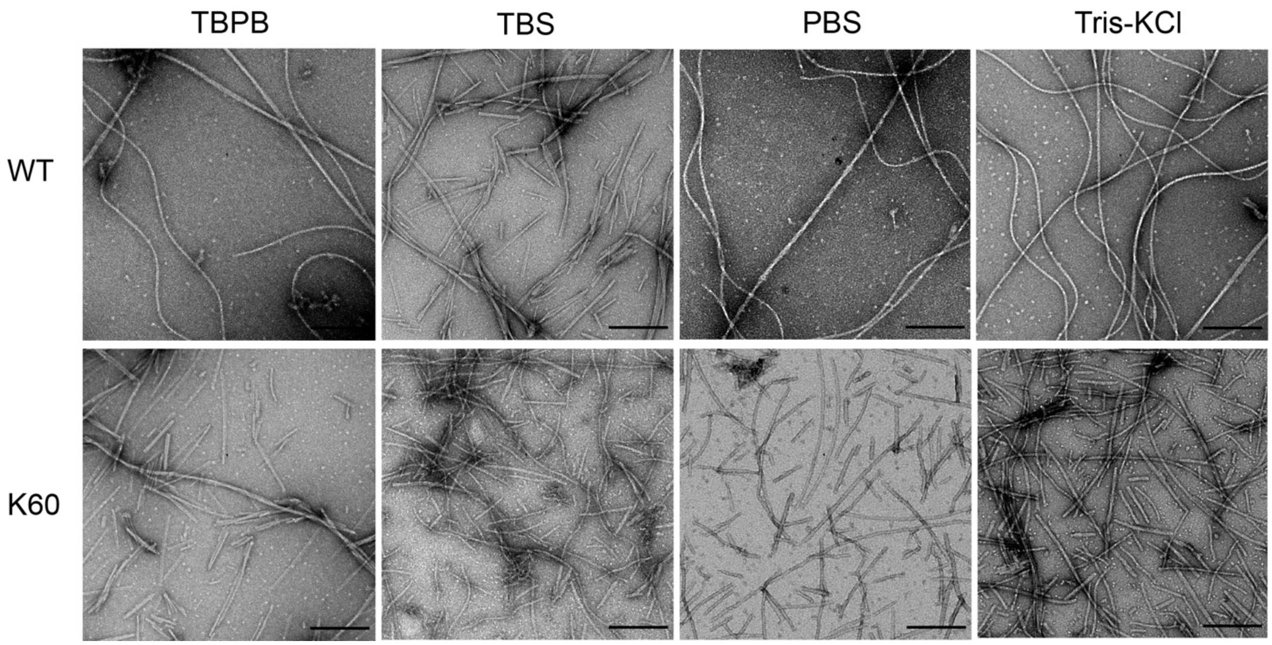


**Figure S10. Wild-type and K60 α-syn fibril buffer screen visualized with negative-stain EM**. TBPB: 15 mM tetrabutylphosphonium bromide, TBS: 20 mM Tris-HCl, 150 mM NaCl pH 8.0, PBS: phosphate buffer saline pH 7.4, Tris-KCl: 20 mM Tris-HCl, 150 mM KCl pH 8.0. Scale bars are 200 nm.

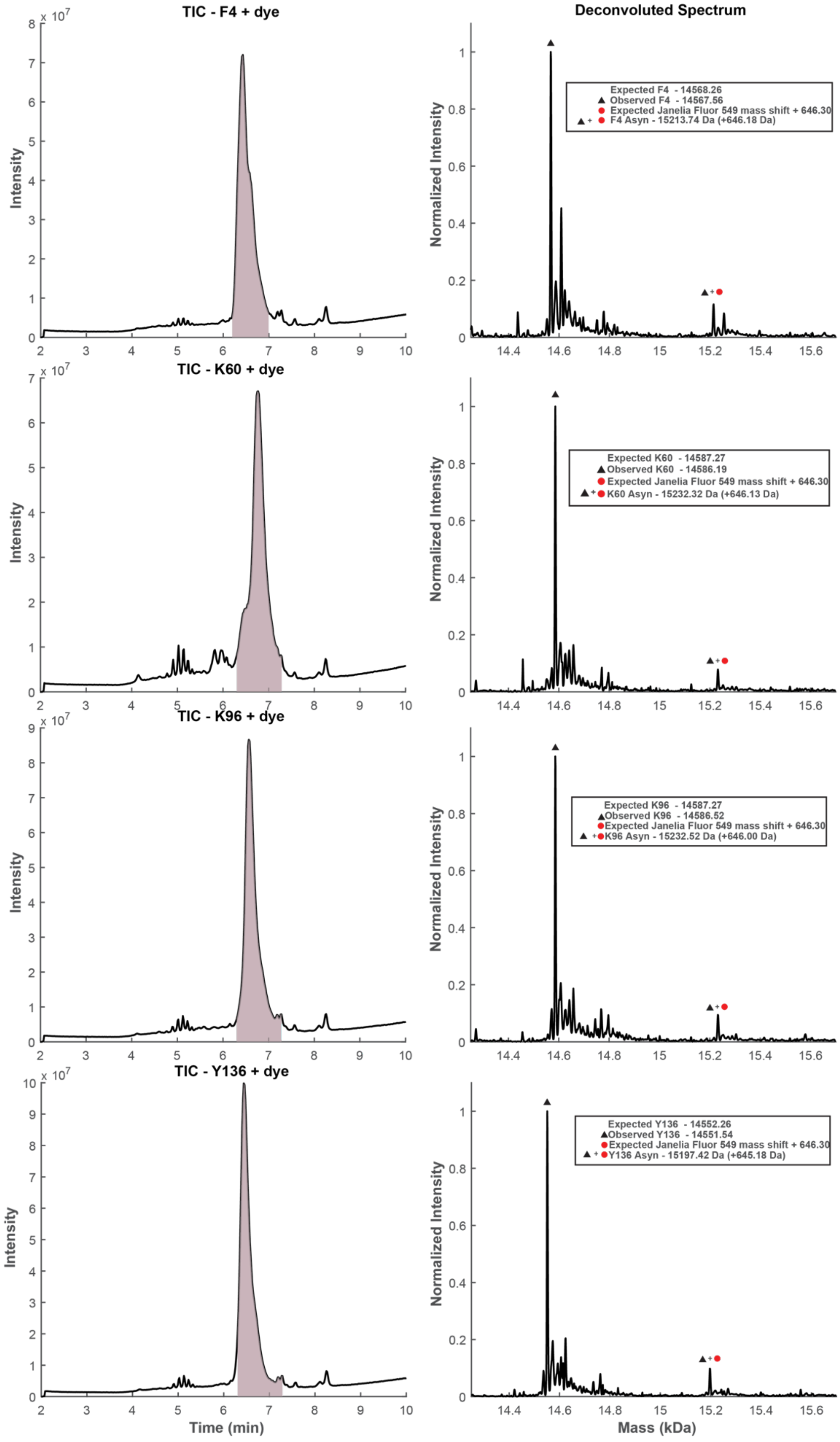
TIC - F4 + dye
Deconvoluted Spectrum
Expected F4 - 14568.26
Observed F4 - 14567.56
Expected Janelia Fluor 549 mass shift + 646.30
F4 Asyn - 15213.74 Da (+646.18 Da)
TIC - K60 + dye
Expected K60 - 14587.27
Observed K60 - 14586.19
Expected Janelia Fluor 549 mass shift + 646.30
K60 Asyn - 15232.32 Da (+646.13 Da)
TIC - K96 + dye
Expected K96 - 14587.27
Observed K96 - 14586.52
Expected Janelia Fluor 549 mass shift + 646.30
K96 Asyn - 15232.52 Da (+646.00 Da)
TIC - Y136 + dye
Expected Y136 - 14552.26
Observed Y136 - 14551.54
Expected Janelia Fluor 549 mass shift + 646.30
Y136 Asyn - 15197.42 Da (+645.18 Da)
Intensity
Normalized Intensity
Time (min)
Mass (kDa)

**Figure S11. LC-MS analysis of Tet2-Et incorporated α-syn monomers clicked with sTCO-JF549 at 1:1 molar ratio.** The total ion chromatogram of each nCAA monomer with the elution peak highlighted in purple indicating the time window averaged for deconvolution (left). The integrated deconvolution mass spectrum from this region, revealing the observed molecular weights for unlabeled protein indicated by the black triangles and observed molecular weights for labeled protein indicated by the black triangles + red circles (right).

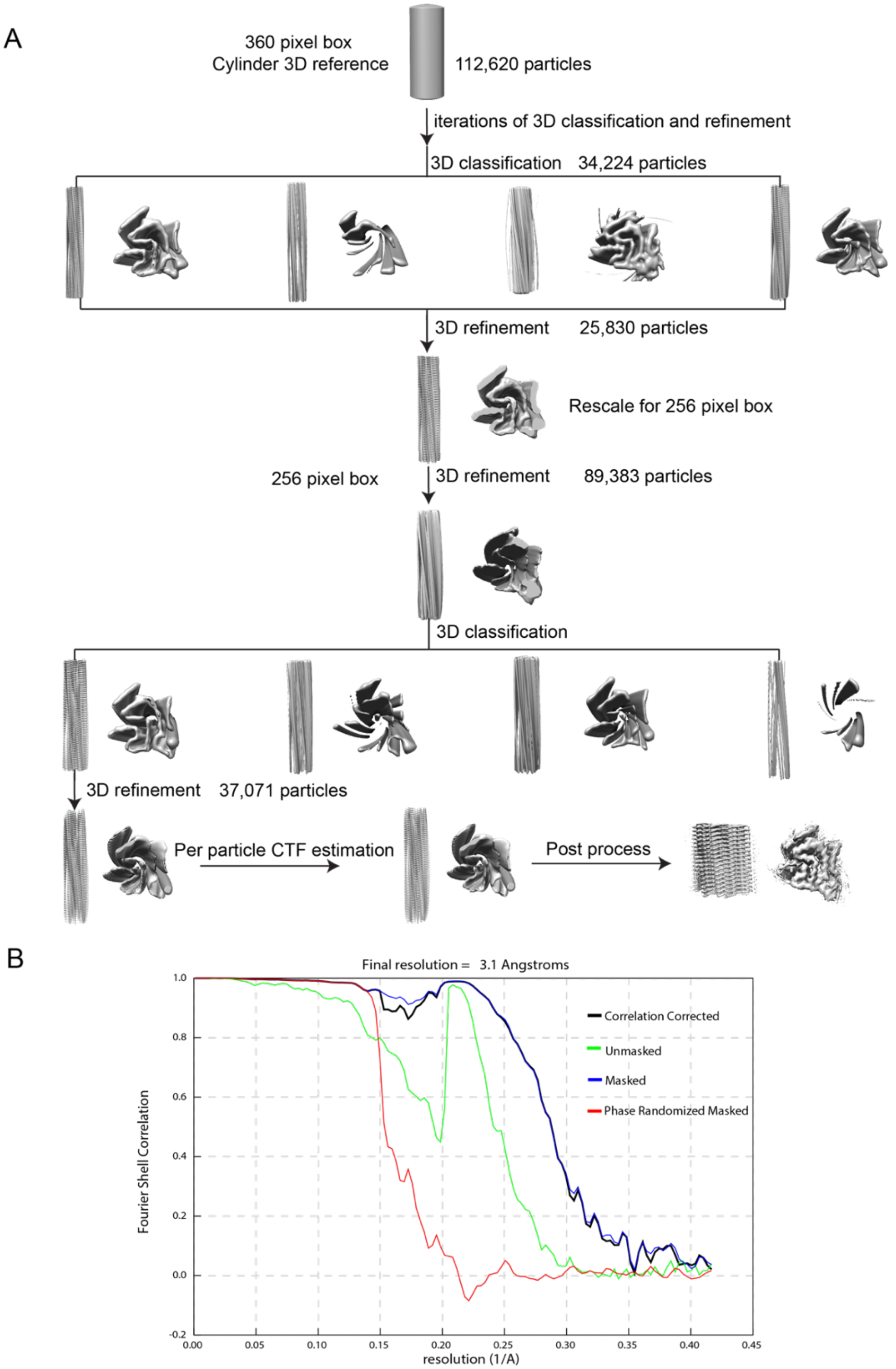


**Figure S12. CryoEM data processing of JF549-Tet2-Et-α-syn fibrils**. **(A)** Schematic representation of helical reconstruction of major twisting species performed in RELION showing fibril maps during 3D refine and 3D classification jobs. **(B)** FSC curves between independently refined half-maps masked and corrected (black), unmasked and corrected (green), masked (navy blue) and phase randomized (red). Final resolution of 3.1 Å was calculated using a cutoff of FSC=0.143.

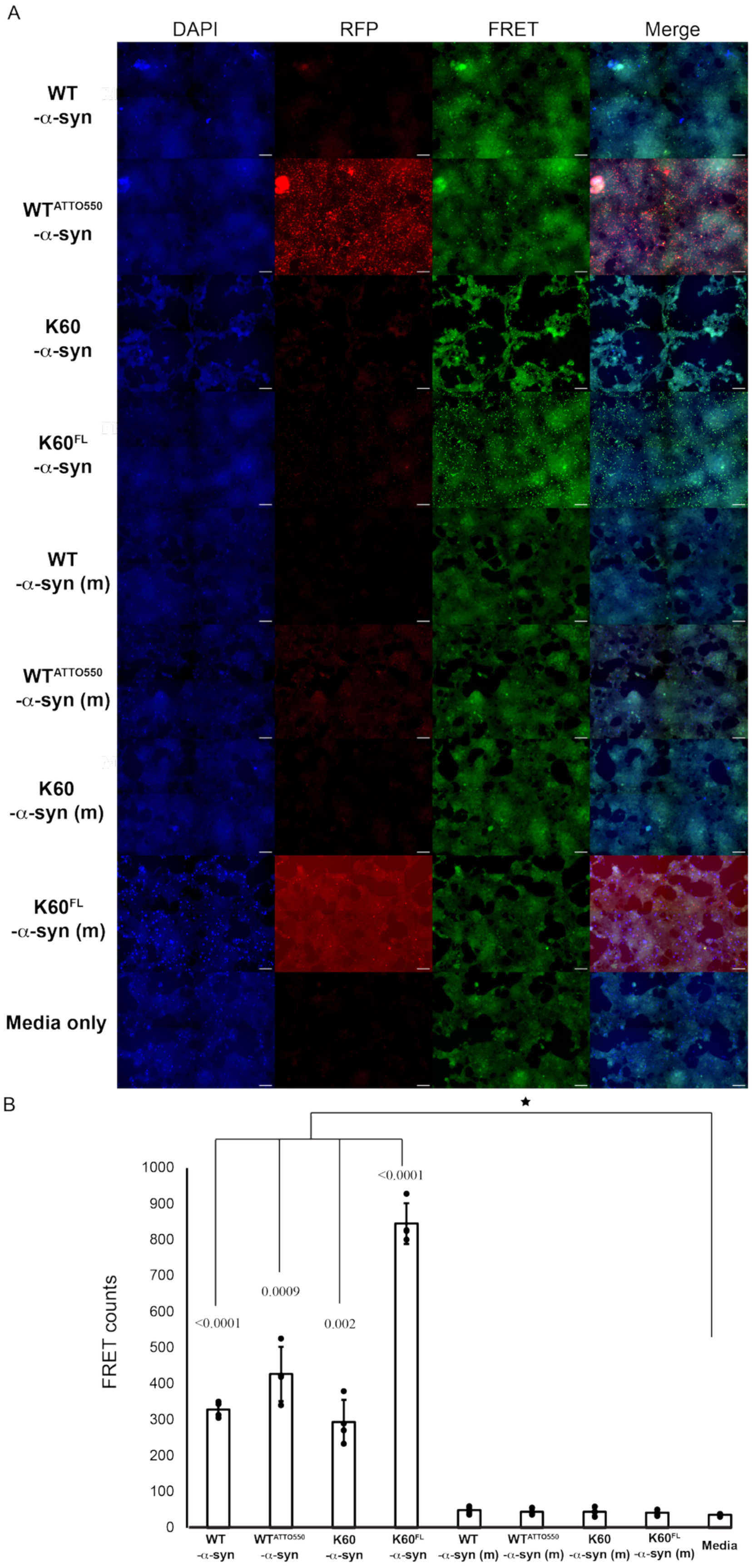

A
DAPI
RFP
FRET
Merge
WT -α-syn
WTATTO550 -α-syn
K60 -α-syn
K60FL -α-syn
WT -α-syn (m)
WTATTO550 -α-syn (m)
K60 -α-syn (m)
K60FL -α-syn (m)
Media only
B
FRET counts
0
100
200
300
400
500
600
700
800
900
1000
<0.0001
0.0009
0.002
<0.0001
★
WT -α-syn
WTATTO550 -α-syn
K60 -α-syn
K60FL -α-syn
WT -α-syn (m)
WTATTO550 -α-syn (m)
K60 -α-syn (m)
K60FL -α-syn (m)
Media

**Figure S13. Uptake and seeding of wild-type and K60 α-syn conjugated ($WT^{ATTO550}$ and $K60^{FL}$) fibrils and monomers in α-syn biosensor cells**. **(A)** 24 hours post-transfection imaged cells with 20x stitching of 9% of the well in a 96 well plate for cells transfected with either labeled (red) or unlabeled α-syn monomers, denoted with (m), or fibrils. Cells stained with NucBlue (blue). Signal is displayed on a linear scale from 900 to 2000 (DAPI), 50 to 700 (RFP) and 200 to 4000 (FRET). Scale bars: 100 μm. **(B)** Quantification of FRET signal in (A). Data are represented as mean total FRET counts for each condition, n = 4 wells . Statistical analysis: Student t test $*p \leq 0.05$ vs. control (media).

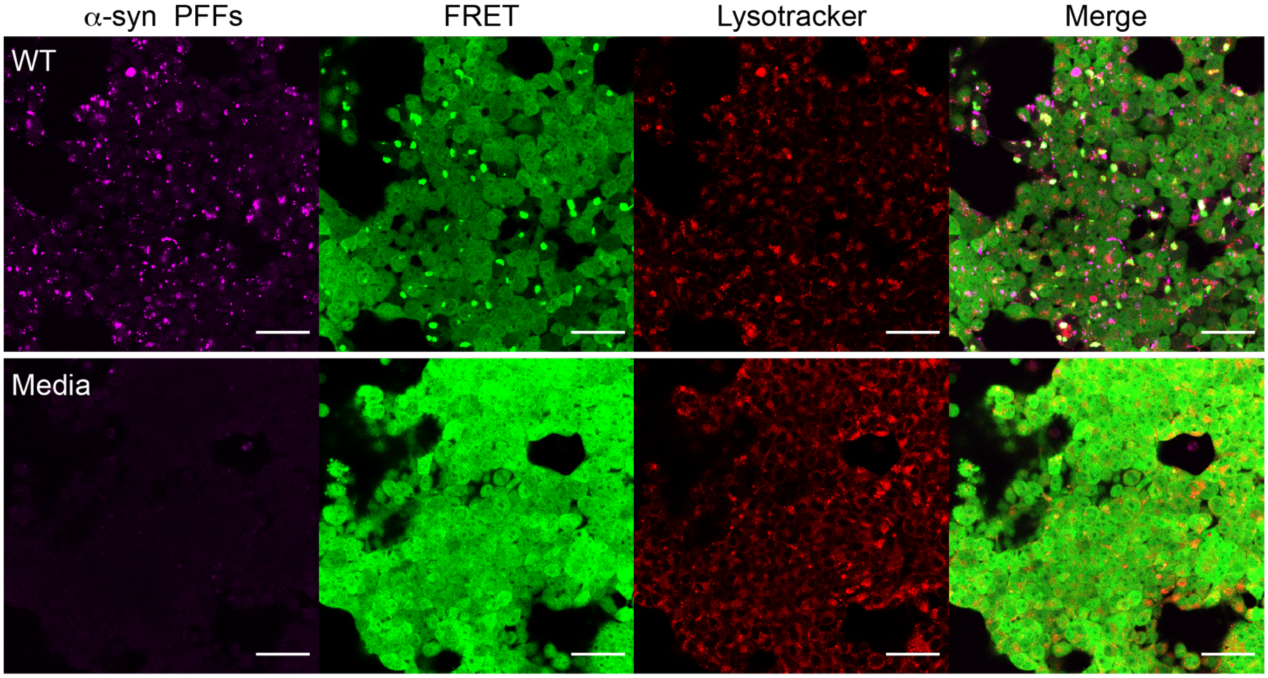


**Figure S14**. **Confocal microscopy of α-syn aggregates in lysotracker-labeled biosensor cells**. 24 hours post-transfection 40x confocal imaging of nascent aggregates, indicated by FRET signal (green), in α-syn biosensor cells seeded with either ATTO-550 wild-type fibril conjugates (magenta) or media only and counterstained with lysotracker deep red (red). Scale bars: 50 µm.

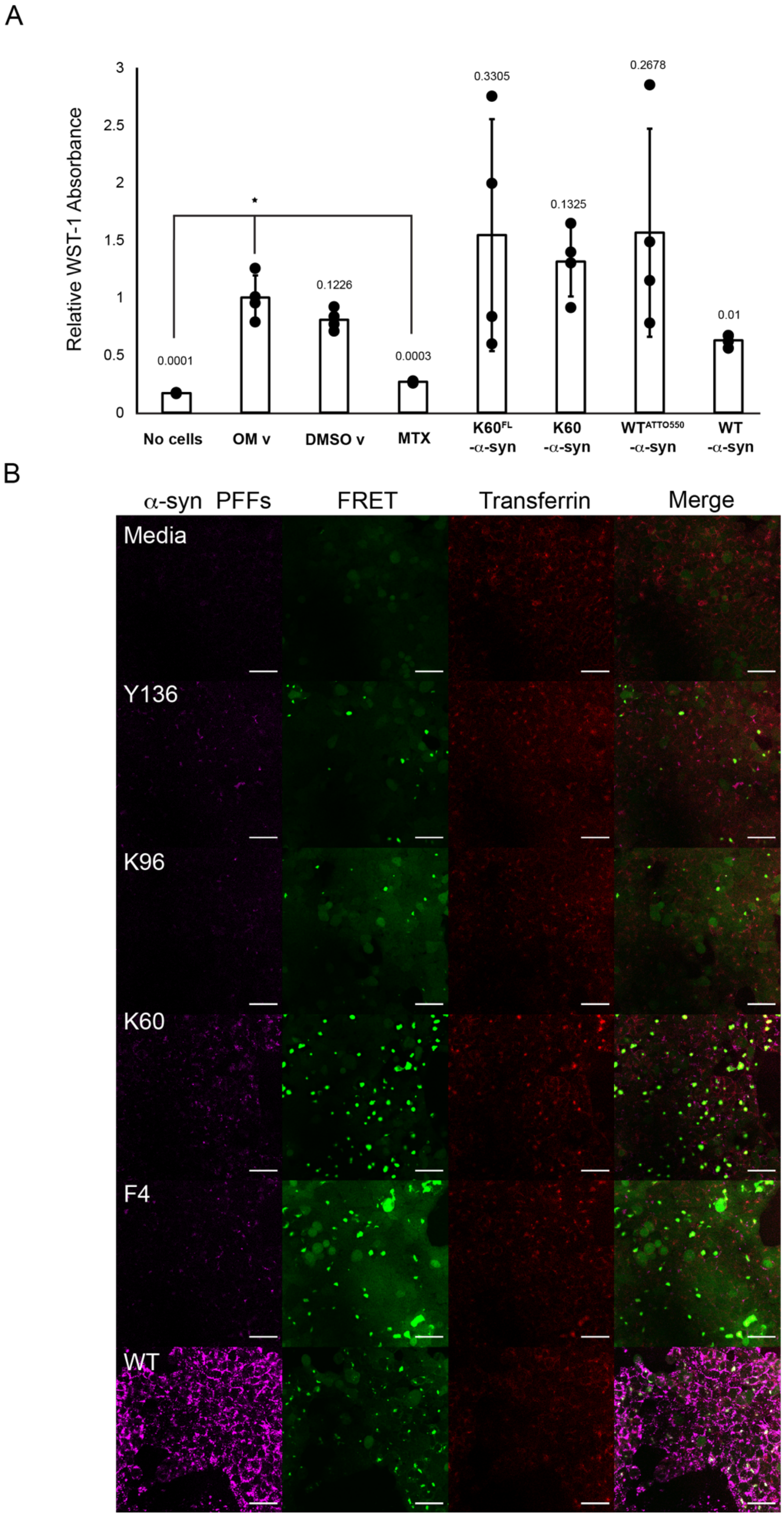

A
Relative WST-1 Absorbance
0
0.5
1
1.5
2
2.5
3
*
0.0001
0.1226
0.0003
0.3305
0.1325
0.2678
0.01
No cells
OM v
DMSO v
MTX
K60FL -α-syn
K60 -α-syn
WTATTO550 -α-syn
WT -α-syn
B
α-syn PFFs
FRET
Transferrin
Merge
Media
Y136
K96
K60
F4
WT

**Figure S15. Functional assays in a-syn biosensor cells treated with fibril conjugates**. **(A)** WST-1 toxicity assay: HEK293T α-syn biosensor cells were grown to 70% confluency and seeded with ATTO-550 wild-type and clicked variant α-syn fibril conjugates. Controls performed included: no cells, Optimem/Lipofectamine 3000 (OM) vehicle, methotrexate (MTX) [5 μM] and DMSO vehicle (0.23% concentration in control wells). 24 hours post-transfection WST-1 reagent was added to cells and measured absorbance at 440 nm. Data are represented as mean absorbance normalized to OM vehicle for each condition, n =4 wells. Statistical analysis: Student t test; $*p < 0.01$ vs. OM vehicle. Error bars = SD. (**B)** HEK293T α-syn biosensor cells were grown to 70% confluency and seeded with ATTO-550 wild-type and clicked variant α-syn fibril conjugates (magenta). 24 hours post-transfection 0.1 mg/ml human transferrin Alexa Fluor 633 (red) was added to cells and nascent aggregates, indicated by FRET signal (green), were visualized by confocal microscopy at 40x and displayed on a linear scale from 0 to 100 (PFFs), and 0 to 100 (Transferrin). Scale bars: 50 μm.

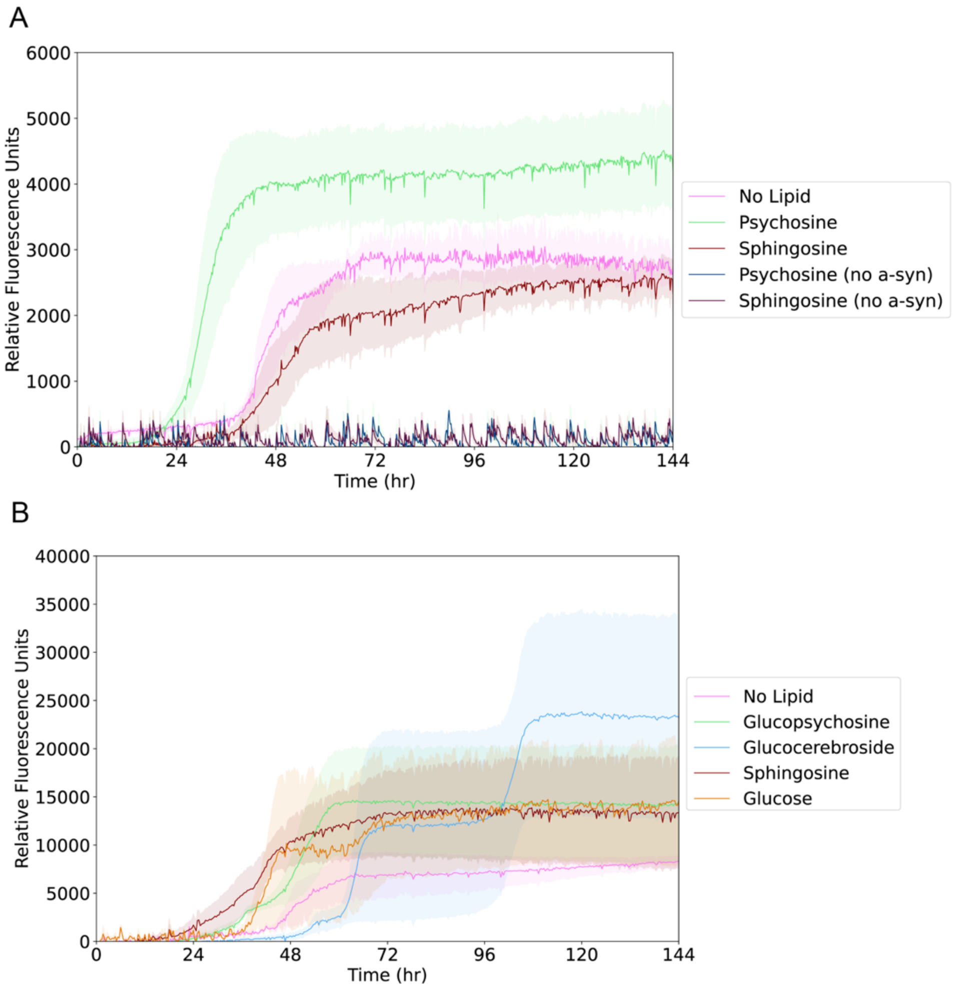


**Figure S16. Thioflavin T assays with α-syn monomers incubated with lipids**. **(A)** Fibrillization assay run with lipid only controls showing lipids psychosine and sphingosine, at 10 µM, do not form ThT-positive aggregates in the absence of α-syn. **(B)** Fibrillization assay with 10 µM lipids related to Gaucher's disease: glucopsychosine.

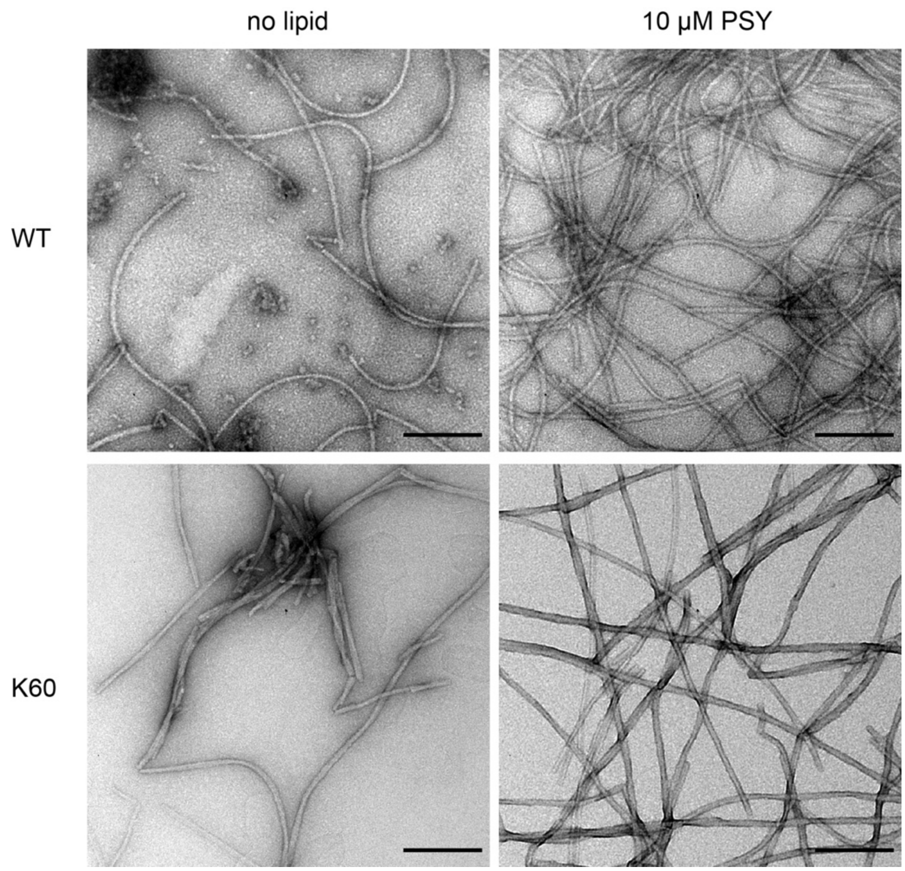


**Figure S17. Negative stain electron microscopy of wild-type and K60 recombinant a-syn fibrils alone or in the presence of psychosine.** Wild-type and K60 α-syn psychosine induced fibrils visualized with negative-stain electron microscopy. Scale bars are 200 nm.

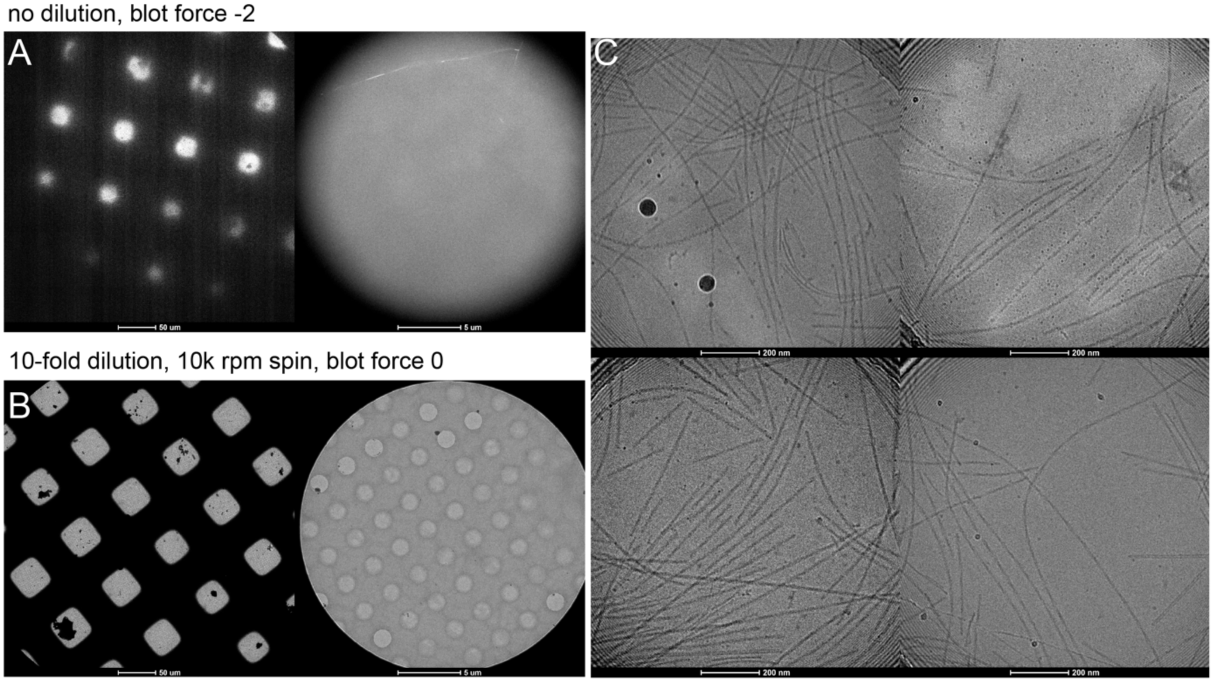


**Figure S18. CryoEM grid optimization of wild-type α-syn psychosine fibrils. (A)** Non-diluted fibrils applied on quantifoil grids are very matted and result in thick ice. **(B)** Combination of a 10-fold dilution of fibrils with a 5 minute spin at 10,000 rpm results in good ice and distribution of fibrils in the holes seen at 57,000x **(C)**. Scale bars 50 μm, 5 μm and 200 nm from left to right.

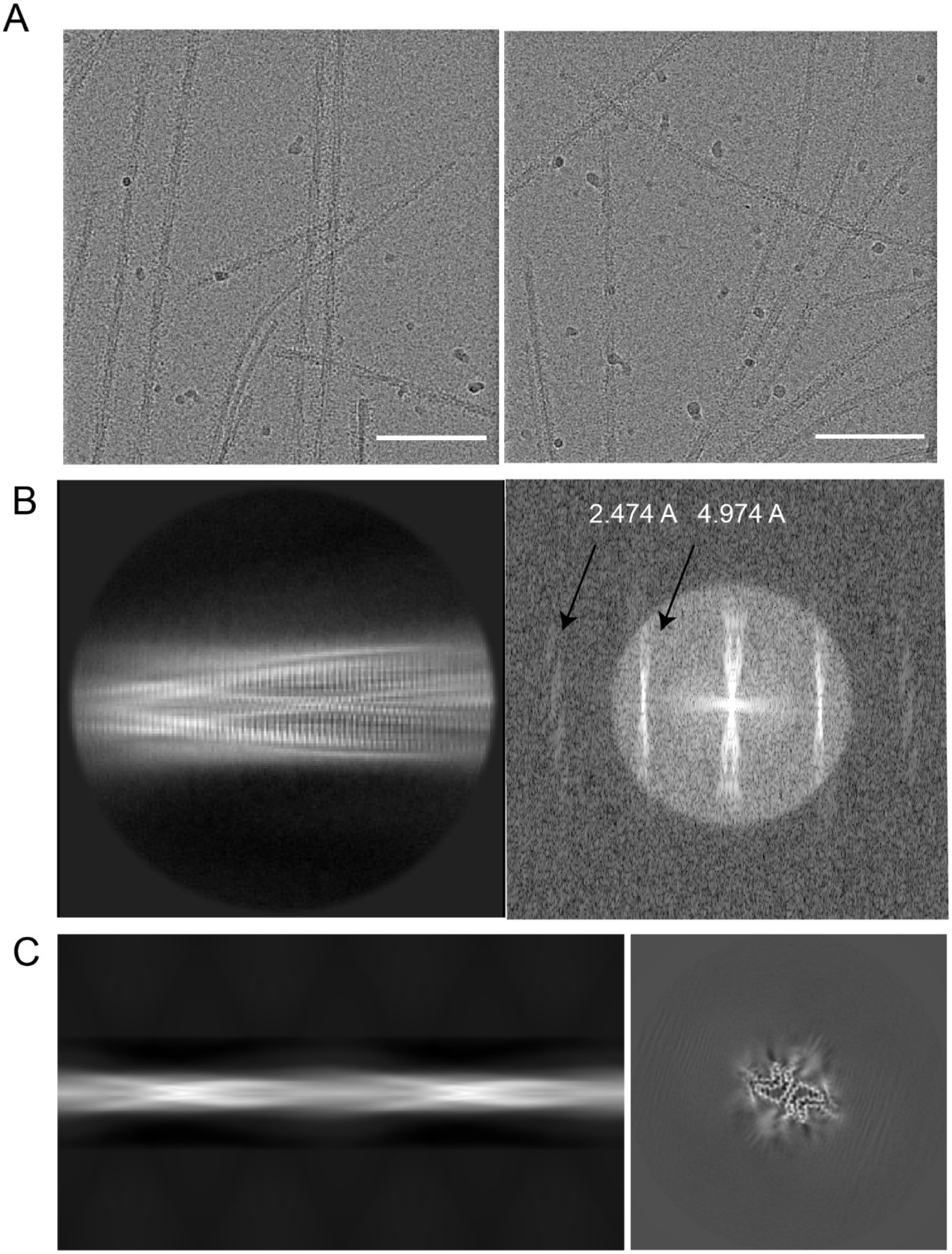


**Figure S19. CryoEM data collection**. **(A)** Representative micrographs during collection displayed on a linear scale from -1000.0 to 1000.0 minimum to maximum brightness and contrast in ImageJ **(B)** Representative 2D class showing beta strand separation. The Fourier transform of this class shows two reflections of 4.974 Å and 2.474 Å, indicating the presence of $P2_1$ symmetry. **(C)** Reprojection of an initial model spanning an entire 750 Å crossover used for the reference in 3D refinement producing the cross-section of the fibril seen on the right.

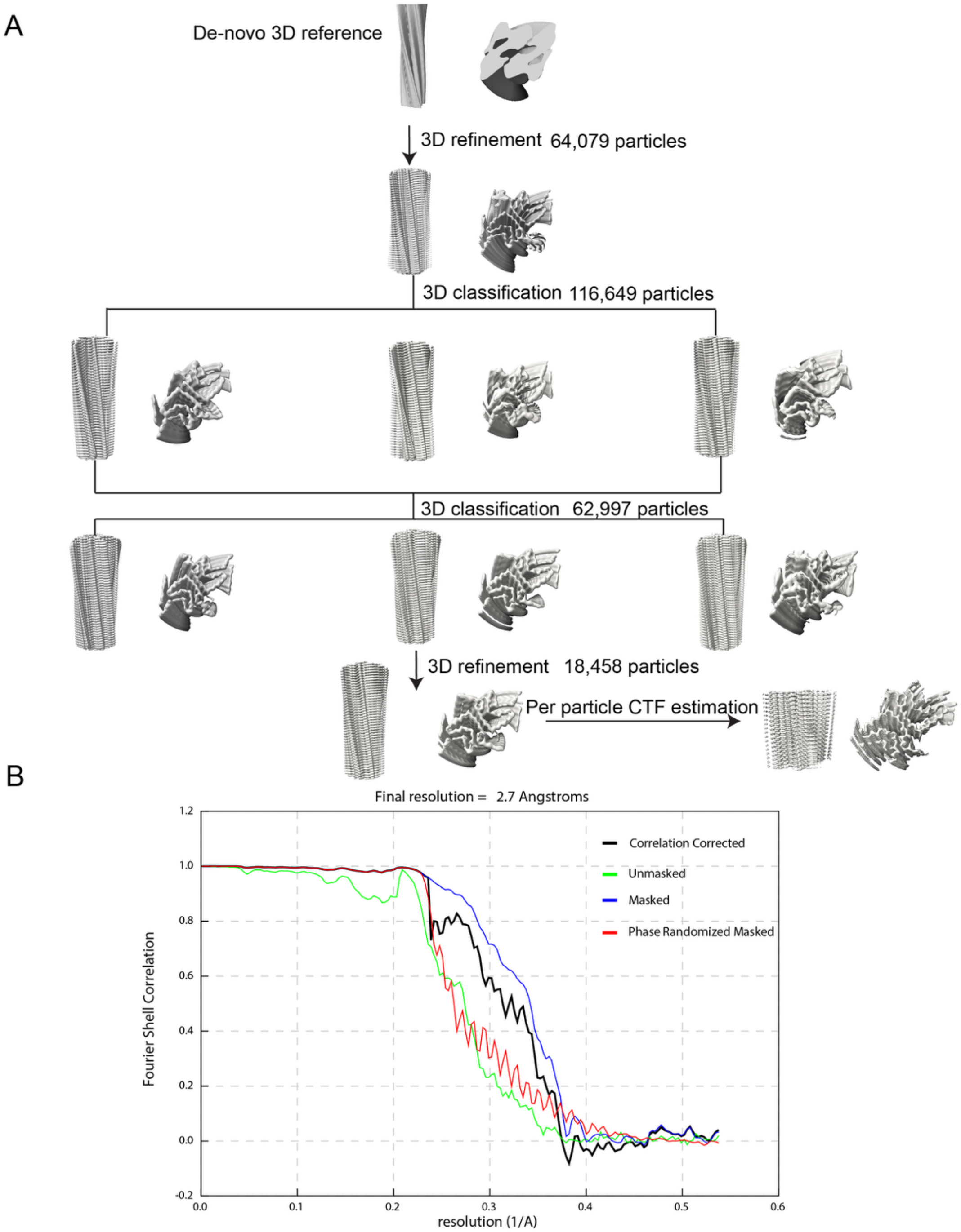


**Figure S20. CryoEM data processing of psychosine-induced α-syn fibrils**. **(A)** Schematic representation of helical reconstruction of major twisting species performed in RELION showing fibril maps during 3D refine and 3D classification jobs. **(B)** FSC curves between independently refined half-maps masked and corrected (black), unmasked and corrected (green), masked (navy blue) and phase randomized (red). Final resolution of 2.7 Å was calculated using a cutoff of FSC=0.143.

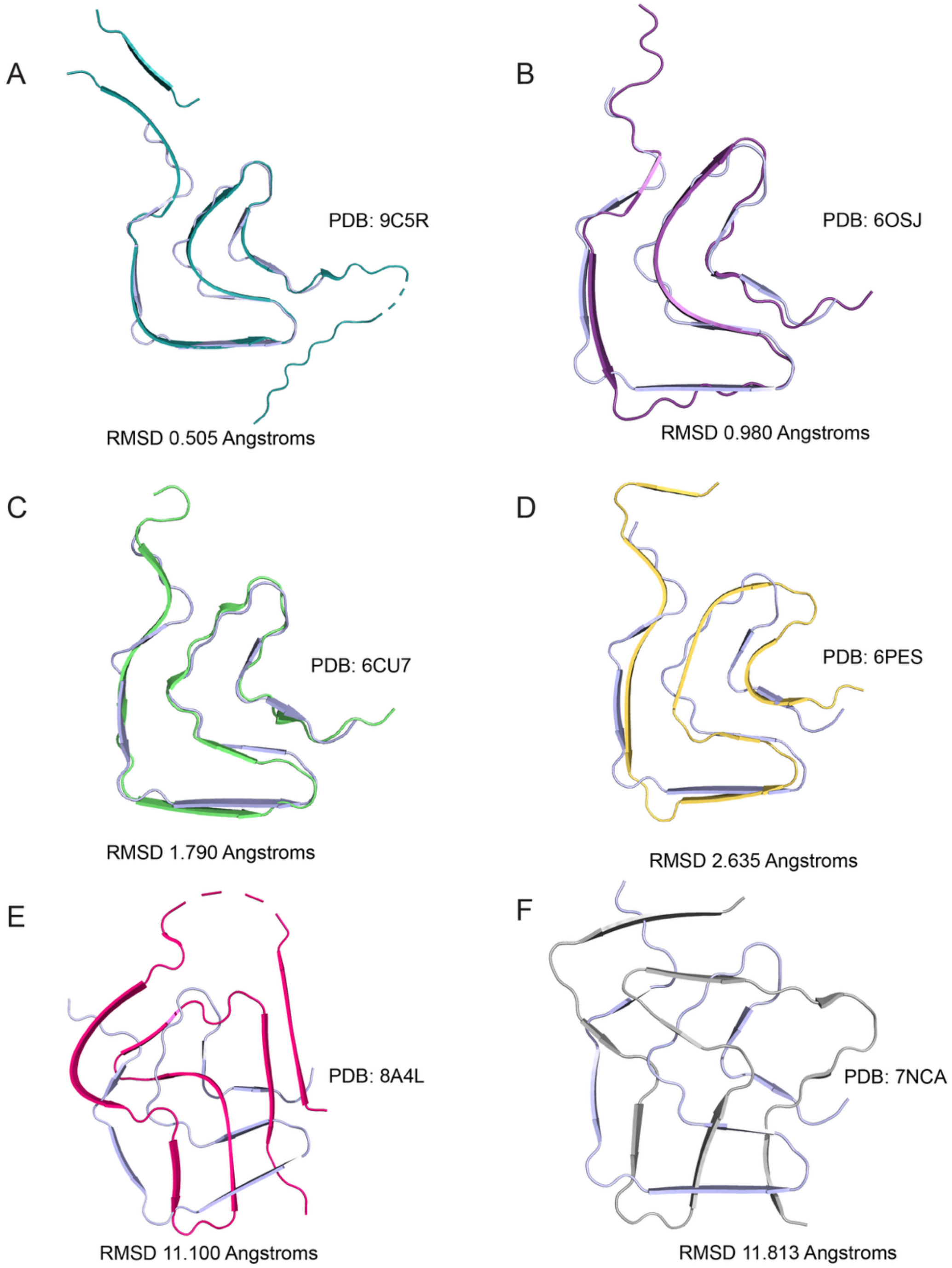


**Figure S21. Structural comparison of psychosine-induced α-syn fibrils to different recombinant α-syn fibrils**. Alignment of this paper's model (light blue) with other recombinant α-syn fibril structures from a seeded full length wild-type fibril (PDB: 9C5R) **(A)**, full-length N-acetylated wild-type (PDB: 6OSJ) **(B)**, full-length wild-type fibril (PDB: 6CU7) **(C)**, full-length H50Q mutated fibril (PDB: 6PES) **(D)**, lipidic fibril (PDB: 8A4L) **(E)**, and a MSA seeded fibril (PDB: 7NCA) **(F)**.

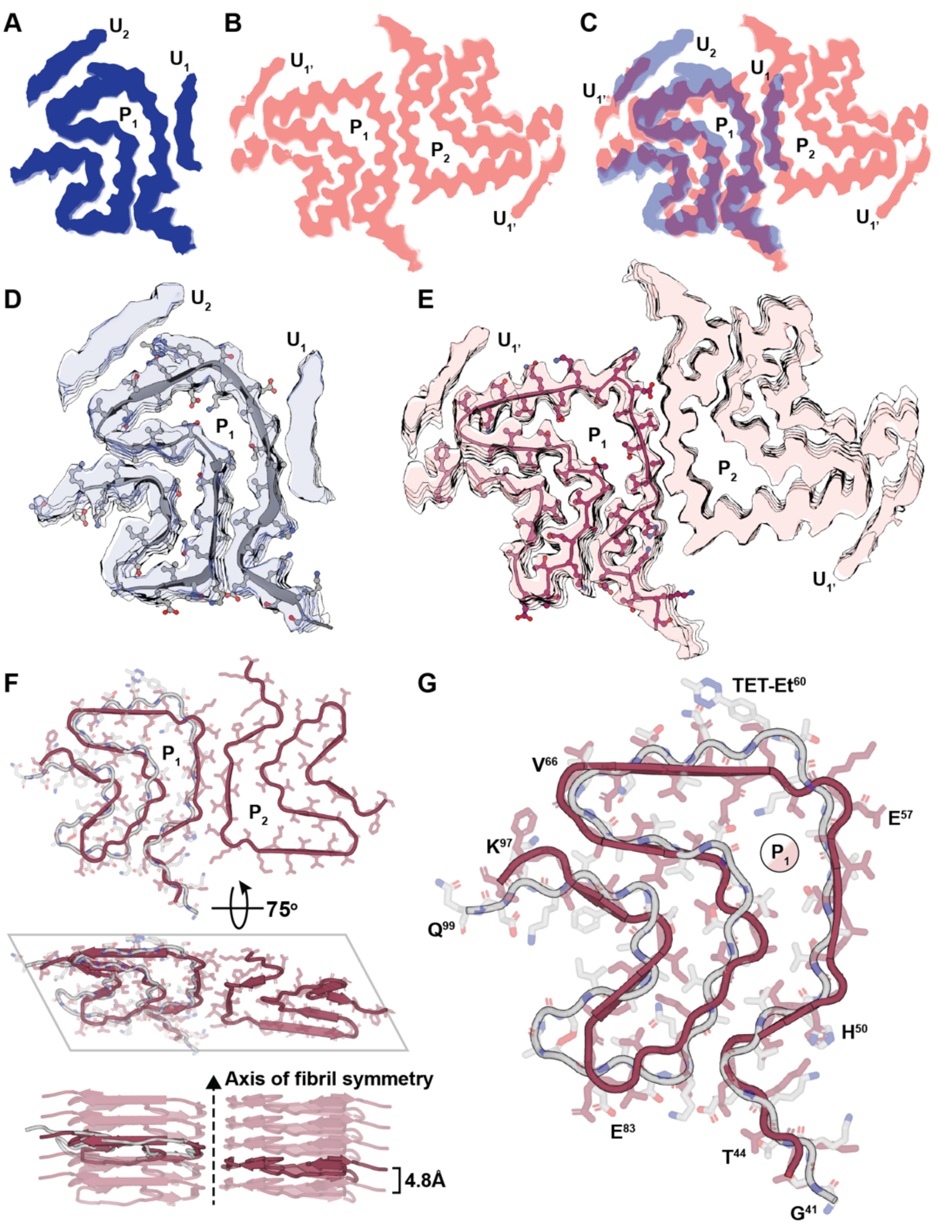


**Figure S22. Comparison of K60$^{FL}$-α-syn and psychosine-induced a-syn fibrils.** A cross section through the core of the K60$^{FL}$-α-syn fibril electrostatic potential map (A, blue) and psychosine-induced α-syn fibril map (B, pink) and their overlay (C). Protofilaments ($P_1$ and $P_2$) in each and isolated unassigned densities $U_1$ and $U_2$ in each are labeled. The overlay in (C) is generated by alignment of the two $P_1$ protofilaments. Models fit into their respective densities are shown for the K60$^{FL}$-α-syn fibril (D) and psychosine-induced fibril (E). An overlay of the models (F) shows their alignment based on $P_1$, viewed down the fibril axis (top), along a 75° tilt angle to the fibril axis (middle), and along the fibril side (bottom). The spacing between layers of the fibril is noted (4.8Å). $P_1$-aligned models enlarged are viewed down the fibril axis, with noteworthy residues labeled, including the TET-Et residue at position 60.
